# An electrodiffusive, ion conserving Pinsky-Rinzel model with homeostatic mechanisms

**DOI:** 10.1101/2020.01.20.912378

**Authors:** Marte J. Sætra, Gaute T. Einevoll, Geir Halnes

**Author notes:** Corresponding author; (GH).

## Abstract

Most neuronal models are based on the assumption that ion concentrations remain constant during the simulated period, and do not account for possible effects of concentration variations on ionic reversal potentials, or of ionic diffusion on electrical potentials. Here, we present what is, to our knowledge, the first multicompartmental neuron model that accounts for electrodiffusive ion concentration dynamics in a way that ensures a biophysically consistent relationship between ion concentrations, electrical charge, and electrical potentials in both the intra- and extracellular space. The model, which we refer to as the electrodiffusive Pinsky-Rinzel (edPR) model, is an expanded version of the two-compartment Pinsky-Rinzel (PR) model of a hippocampal CA3 neuron, where we have included homeostatic mechanisms and ion-specific leakage currents. Whereas the main dynamical variable in the original PR model is the transmembrane potential, the edPR model in addition keeps track of all ion concentrations (Na^+^, K^+^, Ca^2+^, and Cl^−^), electrical potentials, and the electrical conductivities in the intra- as well as extracellular space. The edPR model reproduces the membrane potential dynamics of the PR model for moderate firing activity, when the homeostatic mechanisms succeed in maintaining ion concentrations close to baseline. For higher activity levels, homeostasis becomes incomplete, and the edPR model diverges from the PR model, as it accounts for changes in neuronal firing properties due to deviations from baseline ion concentrations. Whereas the focus of this work is to present and analyze the edPR model, we envision that it will become useful for the field in two main ways. Firstly, as it relaxes a set of commonly made modeling assumptions, the edPR model can be used to test the validity of these assumptions under various firing conditions, as we show here for a few selected cases. Secondly, the edPR model is a supplement to the PR model and should replace it in simulations of scenarios in which ion concentrations vary over time. As it is applicable to conditions with failed homeostasis, the edPR model opens up for simulating a range of pathological conditions, such as spreading depression or epilepsy.

**Author summary:** Neurons generate their electrical signals by letting ions pass through their membranes. Despite this fact, most models of neurons apply the simplifying assumption that ion concentrations remain effectively constant during neural activity. This assumption is often quite good, as neurons contain a set of homeostatic mechanisms that make sure that ion concentrations vary quite little under normal circumstances. However, under some conditions, these mechanisms can fail, and ion concentrations can vary quite dramatically. Standard models are thus not able to simulate such conditions. Here, we present what to our knowledge is the first multicompartmental neuron model that in a biophysically consistent way does account for the effects of ion concentration variations. We here use the model to explore under which activity conditions the ion concentration variations become important for predicting the neurodynamics. We expect the model to be of great use for simulating a range of pathological conditions, such as spreading depression or epilepsy, which are associated with large changes in extracellular ion concentrations.

## Introduction

The neuronal action potential (AP) is generated by a transmembrane influx of Na^+^, which depolarizes the neuron, followed by an efflux of K^+^, which repolarizes it. Likewise, all neurodynamics is fundamentally about the movement of ions, which are the charge carriers in the brain. Therefore, it might seem peculiar that most models of neuronal activity are based on the approximation that the concentrations of the main charge carriers (Na^+^, K^+^, and Cl^−^) do not change over time. This approximation is, for example, incorporated in the celebrated Hodgkin-Huxley model [1], and a large number of later models based on a Hodgkin-Huxley type formalism (see, e.g., [2–7]).

Setting the ion concentrations to not change over time is often a fairly good approximation. The reason is that the number of ions that need to cross the membrane to charge up the neuron by, say, an AP worth of millivolts, is too small to have any notable impact on ion concentrations on either side of the membrane (see, e.g., Box 2.2 in [8]), meaning that concentration changes on a short time scale can be neglected. On a longer time-scale, the ionic exchange due to APs (or other neuronal events), is normally reversed by a set of homeostatic mechanisms such as ion pumps and cotransporters, which work to maintain constant baseline concentrations. In Hodgkin-Huxley type models, the large number of ion pumps, cotransporters and passive ionic leakages that strive towards maintaining baseline conditions are therefore not explicitly modeled. Instead, they are simply assumed to do their job and are grouped into a single *passive* and non-specific leakage current *I*_leak_ = *g*_leak_(*ϕ*_m_ − *E*_leak_), which determines the cell’s resting potential (for a critical study of this approximation, see [9]).

Another approximation commonly applied by modelers of neurons is that the extracellular potential is constant and grounded (*ϕ*_e_ = 0) so that the only voltage variable that one needs to worry about when simulating neurodynamics is the transmembrane potential (*ϕ*_m_). This assumption is implicit in the majority of morphologically explicit models of neurons, where the (spatial) signal propagation in dendrites and axons are computed using the cable equation (see, e.g., [10–12]). Cable-equation based, multicompartmental neuronal models are widely used within the field of neuroscience, both for understanding dendritic integration and neuronal response properties at the single neuron level (see, e.g., [3, 4, 6, 7]) and for exploring the dynamics of large neuronal networks (see e.g., [13–15]). They are even used in the context of performing forward modeling of extracellular potentials, such as local field potentials (LFP), the electrocorticogram (ECoG), and electroencephalogram (EEG) (see, e.g., [16–18]), despite the evident inconsistency involved when first computing neurodynamics under the approximation that *ϕ*_e_ = 0 (Fig 1A), and then in the next step using this dynamics to predict a nonzero *ϕ*_e_ (Fig 1B). The approximation is nevertheless useful since *ϕ*_e_ is typically so much smaller than *ϕ*_m_ that the (ephaptic) effect of *ϕ*_e_ on neurodynamics can be neglected without severe loss in accuracy [19].

**Figure 1.**
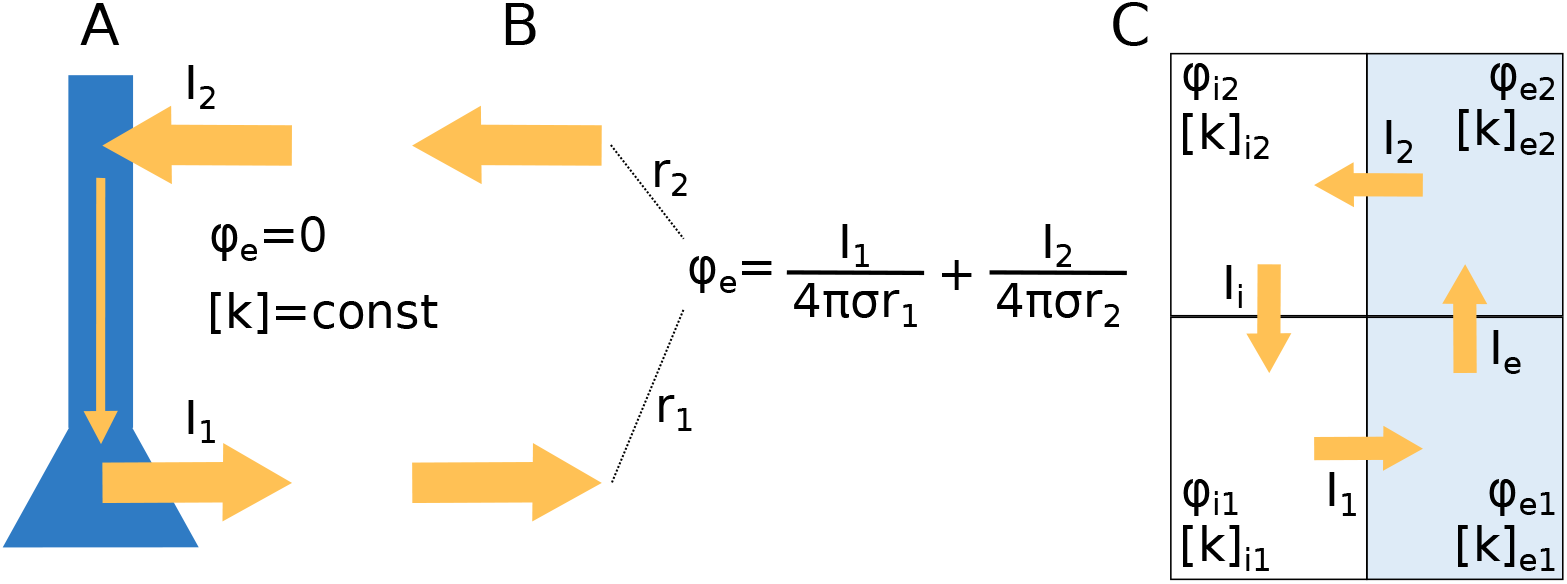
Modeling intra- and extracellular dynamics: standard theory vs. unified framework. **(A)** The dynamics of the membrane potential (*ϕ*_m_) and transmembrane currents of neurons are typically modeled using cable theory. It is then assumed that the extracellular environment is grounded (*ϕ*_e_ = 0). Typically, it is also assumed that ion concentrations both in the intra- and extracellular space are constant, so that also ionic reversal potentials remain constant. **(B)** When knowing the transmembrane neuronal currents (as computed in **(A)**), standard volume conductor theory [20, 21] allows us to estimate the extracellular potential, which is computed as the sum of neuronal point-current sources weighted by their distance to the recording location. An underlying assumption is that fluctuations in *ϕ*_e_ (as computed in **(B)**) are so small that they have no effect on the neurodynamics (as computed in **(A)**), i.e., there is no ephaptic coupling. Another underlying assumption (cf. constant ion concentrations) is that extracellular diffusive currents do not affect electrical potentials. **(C)** We propose a unified, electrodiffusive framework for intra- and extracellular ion concentration and voltage dynamics, assuring a consistent relationship between ion concentrations, electrical charge, and electrical potential in all compartments.

There are, however, scenarios where the assumptions of constant ion concentrations and a grounded extracellular space are not justifiable. Notably, large-scale extracellular ion concentration changes are a trademark of several pathological conditions, including epilepsy and spreading depression [22–25]. In these cases, neurons are unable to maintain their baseline conditions because they for various reasons are too active and/or their homeostatic mechanisms are too slow. During spreading depression, the extracellular K^+^ concentration can change from a baseline value of about 3-5 mM to pathological levels of several tens of mM, and the increased K^+^ concentration tends to coincide with a slow, direct-current (DC) like drop in the extracellular potential, which may be several tens of millivolts in amplitude [25, 26], and can give rise to large spatial gradients. For example, one experiment saw the extracellular K^+^-concentration and *ϕ*_e_ vary by as much as 30 mM and 20 mV, respectively, over the hippocampal depth [26]. Such dramatic gradients in the extracellular environment are likely to have a strong impact on the dynamical properties of neurons, both through the concentration-dependent changes in ion-channel reversal potentials [27–29] and putatively through a direct ephaptic effect from *ϕ_e_* on the membrane potential.

The construction of accurate neuron models that include ion concentration dynamics (and conservation) poses two key challenges. Firstly, ion conserving models need a finely adjusted balance between the homeostatic machinery and all passive and active ion-specific currents so that all ion concentrations, as well as voltages, vary in a biophysically realistic way over time when the neuron is active. Secondly, in spatially extended models, ions will not move only across membranes, but also within the extracellular and intracellular space. Such ionic movement may be propelled both by diffusion and electrical drift. Ionic diffusion can, in principle, affect the electrical potential (since ions carry charge), and the electrical potential can, in principle, affect ion concentration dynamics (since ions drift along potential gradients) [30–32]. Accurate modeling of such systems thus requires a unified, electrodiffusive framework that ensures a physically consistent relationship between ion concentrations, charge density, and electrical potentials.

Intra- or extracellular electrodiffusion is not an issue in single-compartment models, of which there are quite a few that incorporate ion concentration dynamics in a more or less consistent way [28, 29, 33–47]. There are also several morphologically explicit models that have included homeostatic machinery and explicitly simulated ion concentration dynamics (see e.g., [27, 48–57]). However, neither of these have accounted for the electrodiffusive coupling between the movement of ions and electrical potentials (see Results section titled Loss in accuracy when neglecting electrodiffusive effects on concentration dynamics). Hence, to our knowledge, no morphologically explicit neuron model has so far been developed that ensures biophysically consistent dynamics in ion concentrations and electrical potentials during long-time activity, although useful mathematical framework for constructing such models are available [58–62].

The goal of this work is to propose what we may refer to as “a minimal neuronal model that has it all”. By “has it all”, we mean that it (1) has a spatial extension, (2) considers both extracellular- and intracellular dynamics, (3) keeps track of all ion concentrations (Na^+^, K^+^, Ca^2+^, and Cl^−^) in all compartments, (4) keeps track of all electrical potentials (*ϕ*_m_, *ϕ*_e_, and *ϕ*_i_ - the latter being the intracellular potential) in all compartments, (5) has differential expression of ion channels in soma versus dendrites, and can fire somatic APs and dendritic calcium spikes, (6) contains the homeostatic machinery that ensures that it maintains a realistic dynamics in *ϕ*_m_ and all ion concentrations during long-time activity, and (7) accounts for transmembrane, intracellular and extracellular ionic movements due to both diffusion and electrical migration, and thus ensures a consistent relationship between ion concentrations and electrical charge. Being based on a unified framework for intra- and extracellular dynamics (Fig 1C), the model thus accounts for possible ephaptic effects from extracellular dynamics, as neglected in standard feedforward models based volume conductor theory (Fig 1A-B). By ‘‘minimal” we simply mean that we reduce the number of spatial compartments to the minimal, which in this case is four, i.e., two neuronal compartments (a soma and a dendrite), plus two extracellular compartments (outside soma and outside dendrite). Technically, the model was constructed by adding homeostatic mechanisms and ion concentration dynamics to an existing model, i.e., the two-compartment Pinsky-Rinzel (PR) model [3], and embedding in it a consistent electrodiffusive framework, i.e., the previously developed Kirchhoff-Nernst-Planck framework [31, 32, 60, 62]. For the remainder of this paper, we refer to our model as the electrodiffusive Pinsky-Rinzel (edPR) model.

The remainder of this article is organized as follows. First, we present the edPR model and illustrate the numerous variables that it can simulate. Next, we show that the edPR model can reproduce the firing properties of the original PR model. By running long-time simulations (several minutes of biological time) on both models, we identify the firing conditions under which the two models maintained a similar firing pattern, and under which conditions concentration effects became important so that dynamics of the edPR-model diverged from the original PR model over time. Finally, we use the electrodiffusive edPR model to explore the validity of some important assumptions commonly made in the field of computational neuroscience, regarding the decoupling of electrical and diffusive signals. We believe that the IPCR model will be of great value for the field of neuroscience, partly because it gives a deepened insight into the balance between neuronal firing and ion homeostasis, partly because it lends itself to explore under which conditions the common modeling assumption of constant ion concentrations is warranted, and most importantly because it opens for more detailed mechanistic studies of pathological conditions associated with large changes in ion concentrations, such as epilepsy and spreading depression [22–25].

## Results

### An electrodiffusive Pinsky-Rinzel model

The here proposed electrodiffusive Pinsky-Rinzel (edPR) model is inspired by the original Pinsky-Rinzel (PR) model [3], which is a two-compartment (soma + dendrite) version of a CA3 hippocampal cell model, initially developed by Traub et al. [2]. In the original PR model, the somatic compartment contains Na^+^, and K^+^ delayed rectifier currents (*I*_Na_ and *I*_K–DR_), while the dendritic compartment contains a voltage-dependent Ca^2+^ current (*I*_Ca_), a voltage-dependent K^+^ afterhyperpolarization current (*I*_K–AHP_), and a Ca^2+^-dependent K^+^ current (*I*_K–C_). In addition, both compartments contain passive leakage currents. Despite its small number of compartments and conductances, the PR model can reproduce a variety of realistic firing patterns when responding to somatic or dendritic stimuli, including somatic APs and dendritic calcium spikes.

In the edPR model, we have adopted all mechanisms from the original PR model. In addition, we have (i) made all ion channels and passive leakage currents ion-specific, (ii) included a 3Na^+^/2K^+^ pump (*I*_pump_), a K^+^/Cl^−^ cotransporter (*I*_KCC2_), a Na^+^/K^+^/2Cl^−^ cotransporter (*I*_NKCC1_), and a Ca^2+^/2Na^+^ exchanger, and (iii) included two extracellular compartments (outside soma + outside dendrite). To compute the dynamics of the edPR, we used an electrodiffusive KNP-framework for consistently computing the voltage- and ion concentration dynamics in the intra- and extracellular compartments [60]. The model is summarized in Fig 2 and described in details in the Methods section.

**Figure 2.**
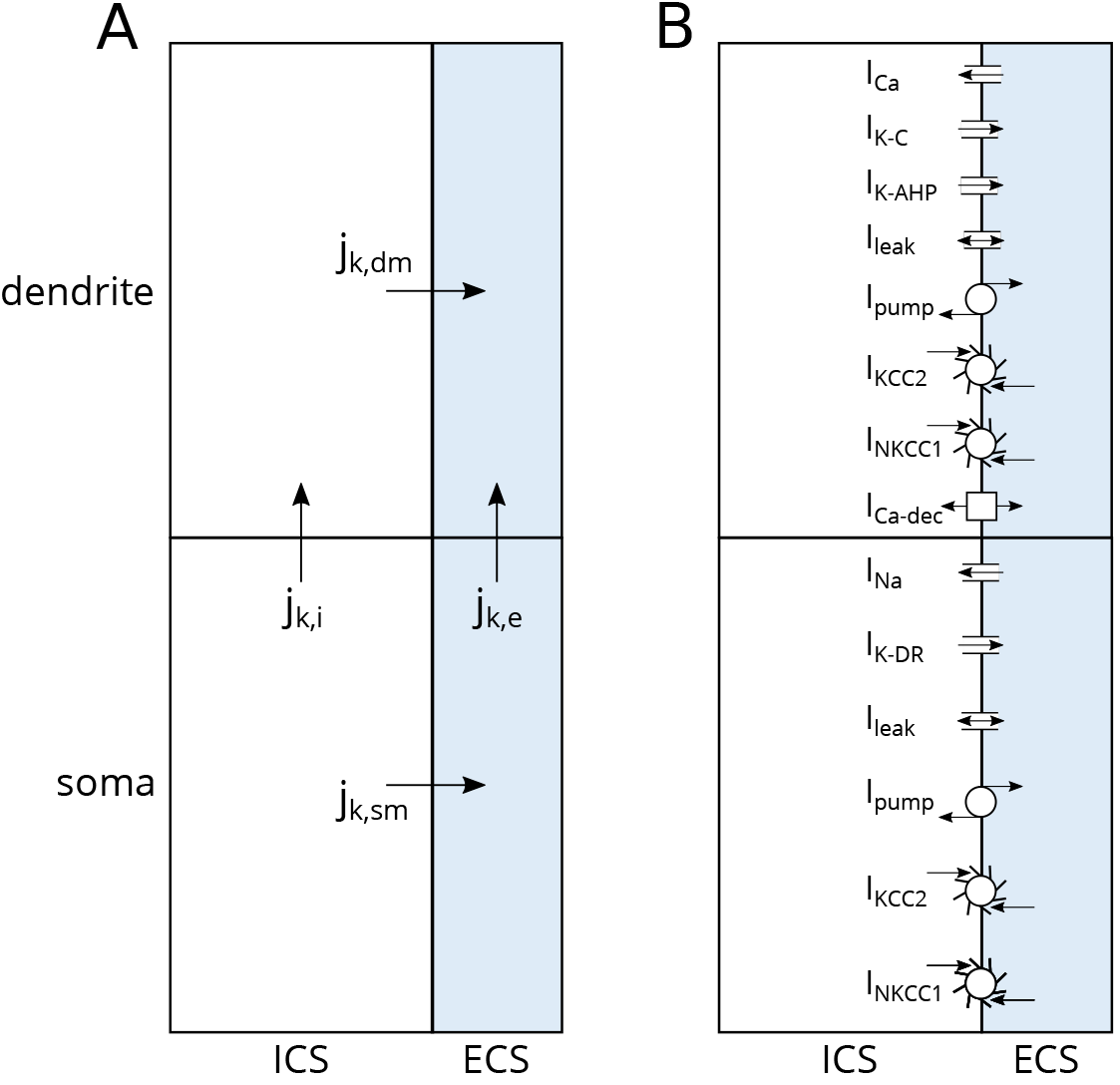
edPR model architecture. **(A)** Two plus two compartments (soma + dendrite), with intracellular space to the left and extracellular space to the right. Two kinds of fluxes of different ion species k are involved: transmembrane fluxes (*j*_k,dm_, *j*_k,sm_) and intra- and extracellular fluxes (*j*_k,i_, *j*_k,e_). The dynamics of the potential *ϕ* and ion concentration dynamics in all compartments were computed using an electrodiffusive framework, ensuring bulk electroneutrality and a consistent relationship between ion concentrations, electrical charge, and voltages. **(B)** Active currents were taken from the original PR model [3]. In the soma, these consisted of Na^+^ and K^+^ delayed rectifier currents (I_Na_ and I_K-DR_). In the dendrite, these consisted of a voltage-dependent Ca^2+^ current (I_Ca_), a Ca^2+^-dependent K^+^ current (I_KC_), and a voltage-dependent K^+^ afterhyperpolarization current (I_K-AHP_). Ion specific passive (leakage-) currents and homeostatic mechanisms were taken from a previous model by Wei et al. [45], and were identical in the soma and dendrite. These included Na^+^, K^+^ and Cl^−^ leak currents, a 3Na^+^/2K^+^ pump (I_pump_), a K^+^/Cl^−^cotransporter (I_KCC2_), and a Na^+^/K^+^/2Cl^−^cotransporter (I_NKCC1_). In addition, the dendrite included a Ca^2+^/2Na^+^ exchanger (I_Ca-dec_), providing an intracellular Ca^2+^ decay similar to that in the PR model.

### Key dynamical variables in the electrodiffusive Pinsky-Rinzel model

While the key variable in the original PR model is the membrane potential *ϕ*_m_, the edPR model allows us to compute a multitude of variables relevant to neurodynamics. The functionality of the edPR model is illustrated in Fig 3, which shows a 60 s simulation where the model fires at 1 Hz for 10 s. We have plotted a selection of output variables, including the membrane potential (Fig 3A-B), extracellular potentials (Fig 3C-D), the dynamics of all ion concentrations in all compartments (Fig 3E-H), concentration effects on ionic reversal potentials (Fig 3I-J), concentration effects on the electrical conductivity of the intra- and extracellular medium (Fig 3K), and ATP consumption (Fig 3L) of the 3Na^+^/2K^+^pump and Ca^2+^/2Na^+^ exchangers.

**Figure 3.**
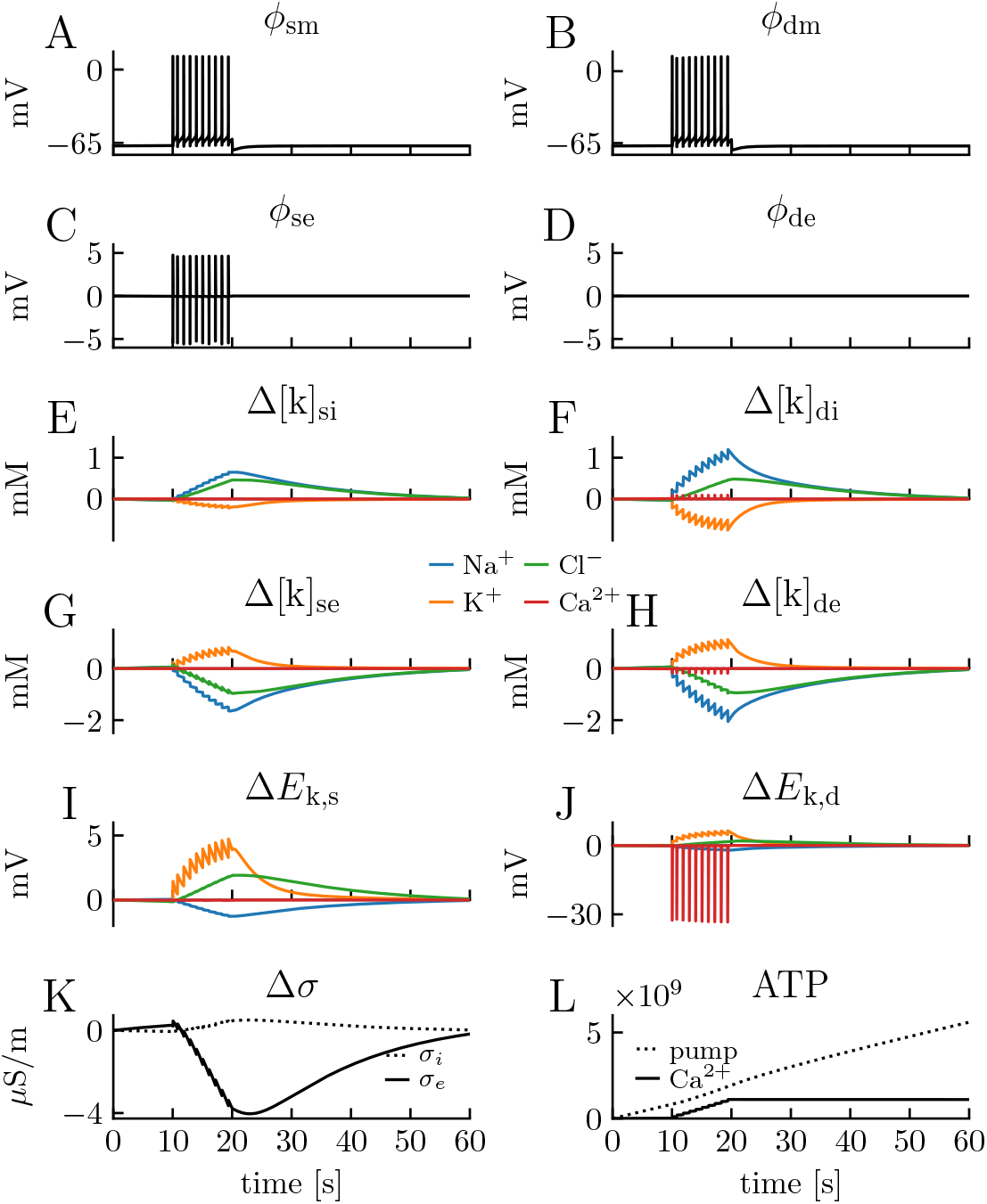
Output of the edPR model. A 28 pA step-current injection was applied to the somatic compartment between *t* = 10 s and *t* = 20 s, and the model responded with a firing rate of 1 Hz. **(A-B)** The membrane potential *ϕ*_m_ of the soma and the dendrite, respectively. **(C-D)** The extracellular (index e) potential *ϕ*_e_ of the soma (index s) and the dendrite (index d), respectively. The dendritic extracellular compartment was chosen as the reference point when calculating potentials, so *ϕ*_de_ was zero by definition. Since amplitudes in *ϕ*_m_ were so much larger than for *ϕ*_e_, intracellular (index i) potentials (*ϕ*_e_ = *ϕ*_e_ + *ϕ*_m_) were similar to *ϕ*_m_, and therefore not shown. **(E-F)** Ion concentrations dynamics of all ion species k (Na^+^Cl^−^, K^+^, Ca^2+^) in all four compartments shown in terms of their deviance from baseline concentrations. **(I-J)** Changes in reversal potentials for all ion species in the soma and the dendrite, respectively. **(K)** Change in conductivity of the intra- and extracellular media (*σ*_i_ and *σ*_e_, respectively). **(L)** Accumulative number of ATP molecules consumed by the 3Na^+^/2K^+^ pumps and Ca^2+^/2Na^+^ exchangers.

Unlike neuronal models based on cable theory, where *ϕ*_e_ is assumed to be zero so that *ϕ*_m_ = *ϕ*_i_, the edPR model computes *ϕ*_m_, *ϕ*_i_, and *ϕ*_e_ from a consistent framework where ephaptic effects from *ϕ*_e_ on *ϕ*_m_ are accounted for (Fig 3C). Due to the electrical coupling between the soma and dendrite, the fluctuations in *ϕ*_m_ were similar in these compartments, and a more detailed analysis of the AP shapes is found further below. While an action potential essentially gave a depolarization followed by a repolarization of *ϕ*_m_, its extracellular signature was essentially a voltage drop (to about −5 mV) followed by a voltage increase (to about +5 mV). This biphasic response of the extracellular AP signature has been seen in several studies (for an analysis, see [20, 21]). In experimental recordings, amplitudes in *ϕ*_e_ fluctuations are typically on the order of 100 *μ*V, which is much smaller than that predicted by the edPR model. The discrepancy is an artifact that is mainly due to the 1D approximation in the edPR-model (see Discussion). The dendritic extracellular potential (Fig 3D) was by definition zero at all times, as this compartment was used as the reference point for the potential.

The effect of neuronal firing on the ion concentration dynamics is illustrated in Fig 3E-H. Before the stimulus onset, the cell was resting at approximately −68 mV, and ion concentrations remained at baseline values. During AP firing, the ion concentrations varied in a jigsaw-like fashion. As the extracellular volume was set to be half as big as the intracellular volume, changes in extracellular ion concentrations were about twice as big as the changes in intracellular ion concentrations. The jigsaw pattern was most pronounced for the K^+^ and Na^+^ concentrations, as these were the main mediators of the APs (Fig 3E-H). The pattern reflects a cycle of (i) incremental steps away from baseline concentrations, which were mediated by the complex of mechanisms active during the APs, followed by (ii) slower decays back towards baseline, which were mediated by pumps and cotransporters working between the APs. In this simulation, the decay was incomplete, so that concentrations reached gradually larger peak values by each consecutive AP. However, as we show later (see Section titled The edPR model predicts homeostatic failure due to high firing rate), the concentrations did, in this case, approach a firing-frequency dependent steady state. When the firing ceased, the pumps and cotransporters could work uninterruptedly to re-establish the baseline ion concentrations. Although a full recovery of ionic concentrations took on the order of 30 s, the resting membrane potential of about-68 mV was recovered much faster (ms timescale), after which the recovery process was due to an electroneutral exchange of ions between the neuron and the extracellular space.

As ion concentrations varied during the simulation, so did the ionic reversal potentials, *E*_k_(Fig 3I-J). The by far largest change was seen for the Ca^2+^ reversal potential in the dendrite (*E*_k,d_), which dropped by as much as −30mV during an AP, (i.e., from a baseline value of 124 mV to 94 mV). The explanation is that the basal intracellular Ca^2+^-concentration is extremely low (100 nM) compared to the concentrations of other ion species (several mM), and therefore experienced a much larger relative change during the simulation. Among the main charge carriers (Na^+^Cl^−^, K^+^), the lowest concentration is found for K^+^in the extracellular space (Table 5 in Methods). For that reason, the second largest change in reversal potential was found for *E*_K_, which increased by about 5 mV (i.e., from a basal value of −84 mV to −79 mV) in both the soma and dendrite. The changes in *E*_Ca_ and *E*_κ_ had a relatively minor impact on the firing pattern in the shown simulations, as the relative change in the driving force *ϕ*_m_ – *E*_k_ was not that severe.

The conductivities (*σ*) of the intra- and extracellular bulk solutions depend on the availability of free charge carriers, and are in the edPR model functions of the ion concentrations and their mobility (cf. Eq 19). The changes in *σ* were minimal during the conditions simulated here (Fig 3K), i.e., *σ* varied by a few *μ*S/m over the course of the simulation, while the basal levels were approximately 0.08 S/m and 0.67 S/m for the intra- and extracellular solutions, respectively.

Finally, the 3Na^+^/2K^+^ pump and Ca^2+^/2Na^+^ exchanger use energy in the form of ATP to move ions against their gradients. The 3Na^+^/2K^+^ pump exchanges 3 Na^+^ ions for 2 K^+^ions, and consumes one ATP per cycle [63], while we assumed that the Ca^2+^/2Na^+^ exchangers consumed 1 ATP per cycle (i.e., per Ca^2+^ exchanged, as in [64]). As the edPR model explicitly models these processes, we could compute the ATP (energy) consumption of the pumps during the simulation. Fig 3L shows the accumulative number of ATP consumed from the onset of the simulation. The 3Na^+^/2K^+^ pump was constantly active, as it combated leakage currents and worked to maintain the baseline concentration even before stimulus onset. Before stimulus onset, it consumed ATP at a constant rate (linear curve), which increased only slightly at *t* = 10 s when the neuron started to fire (small dent in the curve). As the neuron did not contain any passive leakage of Ca^2+^, the Ca^2+^/2Na^+^ exchangers were only active while the neuron was firing. During firing, the Ca^2+^/2Na^+^ exchanger combated the Ca^2+^entering through the dendritic Ca^2+^ channels, and then consumed approximately the same amount of energy as the 3Na^+^/2K^+^ pump (parallel curves). A high metabolic cost of dendritic Ca^2+^ spikes has previously been reported also in cortical layer 5 pyramidal neurons [64].

We note that the edPR model had a stable resting state before stimulus onset and that it returned to this resting state after the stimulus had been turned off. In this resting state, ion concentrations remained constant, and *ϕ*_m_ was approximately −68 mV. This resting equilibrium was due to a balance between the ion-specific leakage channels, pumps, and cotransporters, which we adopted from previous studies (see Methods). However, the existence of such a homeostatic equilibrium was not highly sensitive to the choice of model parameters. As we confirmed through a sensitivity analysis, varying membrane parameters with ± 15% of their default values, only lead to a variation of about ± 2 mV in the resting potential, see Fig 4.

**Figure 4.**
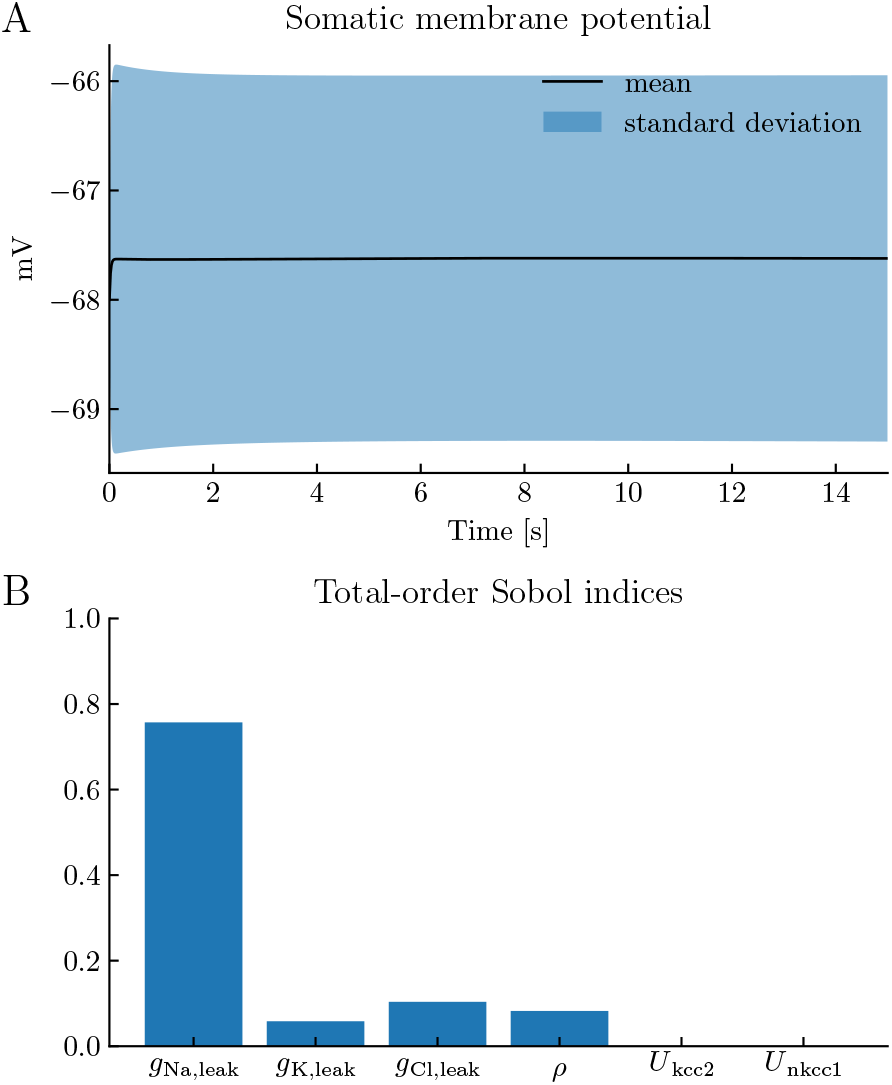
Sensitivity analysis. To study the steady state’s sensitivity to the values of the leak conductances *ḡ*_Na,leak_,*ḡ*_K,leak_, and *ḡ*_C1,leak_, the pump strength *ρ*, and the cotransporter strengths *U*_nkcc1_ and *U*_kcc2_, we performed a sensitivity analysis using Uncertainpy, a Python toolbox for uncertainty quantifications and sensitivity analysis [65]. We ran the model for 15 seconds and let the parameters have a uniform distribution within a ±15% interval around their default values. **(A)** The mean and standard deviation of the somatic membrane potential. We see that the homeostatic equilibrium of the model was not highly sensitive to the choice of model parameters. **(B)** The total-order Sobol indices for the different parameters. We see that the relatively small variation in the potential was mostly due to the variation of *ḡ*_Na,leak_. This makes sense, knowing that the sodium reversal potential (55 mV) is furthest away from the resting potential (≈ −68 mV), making the driving force (*ϕ*_m_ − *E*_k_) of the Na^+^ leak current stronger than those for the other ion-specific leak currents.

### The edPR model reproduces the short term firing properties of the original PR model

A motivation behind basing the electrodiffusive (edPR) model on a previously developed (PR) model, was that we wanted to use the firing properties of the original PR model as a “ground truth” when constraining the edPR model. In particular, we wanted the edPR model to qualitatively reproduce the interplay between somatic action potentials and dendritic Ca^2+^ spikes, as this was an essential feature of the original PR-model [3]. In the PR model, this interplay depended strongly on the coupling strength (coupling conductance) between the soma and dendrite compartment. A weak coupling resulted in a wobbly ping-pong effect, where a somatic AP triggered a dendritic Ca^2+^ spike, which in turn fed back to the soma, giving rise to secondary oscillations in *ϕ*_m_ (Fig 5A). With a strong (about five times stronger) coupling, the somatic and dendritic responses became more similar in shape, as expected (Fig 5B).

**Figure 5.**
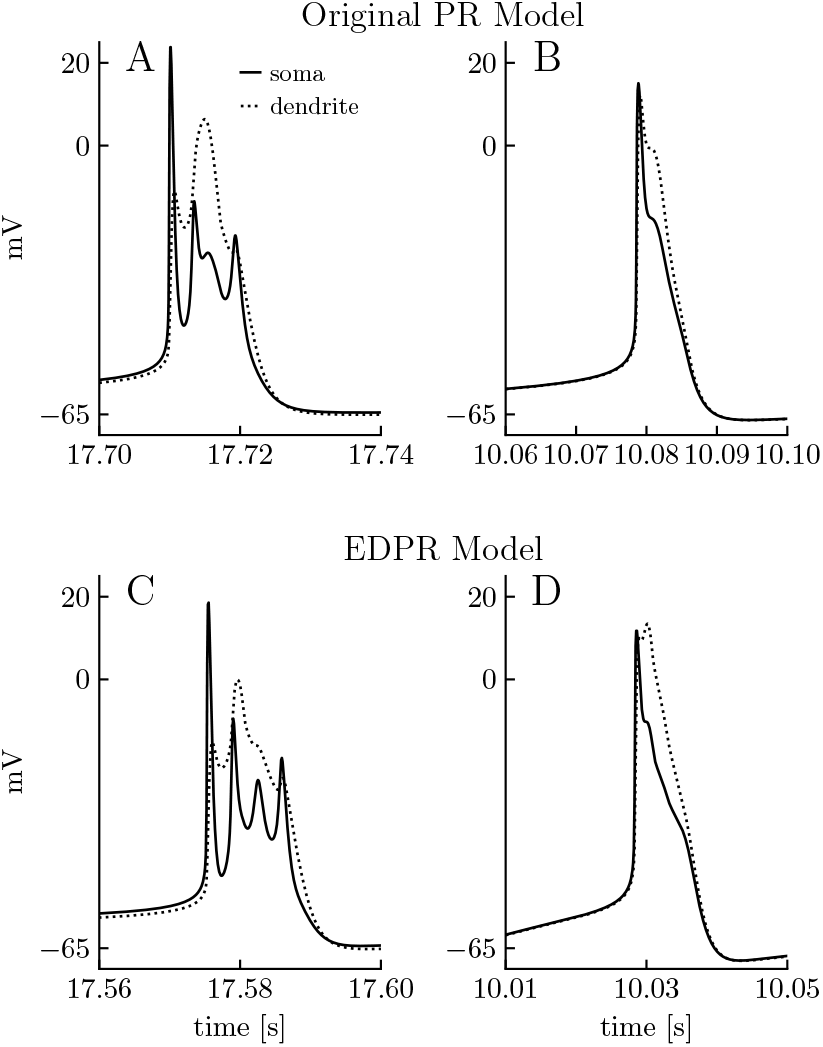
Short term dynamics of the PR and edPR models. The original PR model (top row) and the edPR model (bottom row) exhibit the same spike shape characteristics. **(A)** Spike shape in PR model for weak coupling (low coupling conductance) between the soma and the dendrite. **(B)** Spike shape in PR model for strong (high intracellular conductivity) coupling between the soma and the dendrite. **(C)** Spike shape in edPR model for weak coupling between the soma and the dendrite. **(D)** Spike shape in edPR model for strong coupling between the soma and the dendrite. **(A-D)** A step-stimulus current was turned on at t = 10s, with stimulus strength being 1.35μA/cm^2^ in **(A)**, 0.78μA/cm^2^ in **(B)**, 31 pA in **(C)**, and 28 pA in **(D)**. The panels show snapshots of a selected spike. See the Parameterizations section in Methods for a full description of the parameters used.

Since the edPR model contained membrane mechanisms and ephaptic effects not present in the PR model, selected parameters in the edPR model had to be re-tuned in order to obtain similar firing as the PR model (see Methods). With the selected parameterization of the edPR model (see the Parameterizations section), we were able to reproduce the characteristic features seen in the PR model for a weak (Fig 5C) and strong (about five times stronger) coupling between the soma and dendrite (Fig 5D).

### The edPR model predicts homeostatic failure due to high firing rate

As previously discussed, the PR model was, as most existing neuronal models, constructed under the assumption that ion concentration effects are negligible, an assumption that is justified for short term neurodynamics, and for long term dynamics provided that the activity level is sufficiently low for the homeostatic mechanisms to maintain concentrations close to baseline over time. Hence, we expect there to be a scenario (S1) with a moderately low firing rate, where the PR and edPR can fire continuously and regularly over a long time exhibiting similar firing properties, and another scenario (S2) with a higher firing rate, where the PR and edPR models exhibit similar firing properties initially in the simulation, but where the dynamics of the two models diverge over time due to homeostatic failure accounted for by the edPR model, but not the PR model (which ad hoc assume perfect homeostasis). Simulations of two such scenarios are shown in Figs 6 and 7.

**Figure 6.**
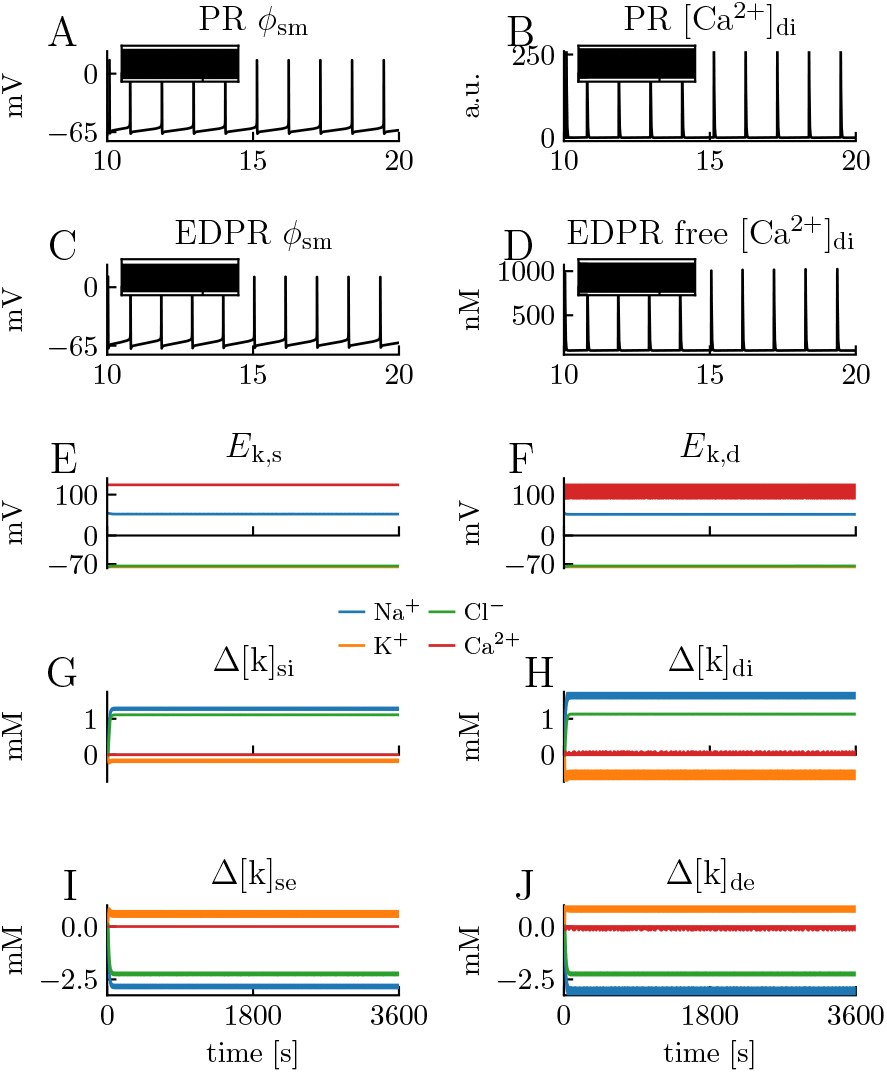
Model comparison for scenario with low frequency firing. Simulations on the PR model and edPR model when both models are driven by a constant input, giving them a firing rate of about 1 Hz. Simulations covered one hour (3600 s) of biological time. **(A-D)** A 10 s sample of the dynamics of the somatic membrane potential *ϕ*_m_ and dendritic Ca^2+^ concentration in the PR model **(A-B)**and edPR model **(C-D)**. This regular firing pattern was sustained over the full 3600 s simulation in both models (inset panels). **(D)** Of the total amount of intracellular Ca^2+^, only 1% (as plotted) was assumed to be free (unbuffered). **(E-F)** Ionic reversal potentials and **(G-J)** ion concentrations in the edPR model did not vary on a long time scale. Indices *i, e, s*, and *d* indicate *intracellular, extracellular, soma*, and *dendrite*, respectively. **(A-J)** Stimulus onset was *t* = 10 s in both models, and stimulus strength was *i*_stim_ = 0.78*μ*A/cm^2^ in the PR model **(A-B)** and *i*_stim_ = 28 pA in the edPR model **(C-J)**. See the Parameterizations section in Methods for a full description of the parameters used.

To simulate scenario S1, the PR model (Fig 6A-B) and edPR model (Fig 6C-J) were given a constant input (see figure caption) that gave them a firing rate of about 1 Hz. Both models then settled at a regular firing rate, and in neither of them the firing pattern changed over time, even in simulations of as much as an hour of biological time. For the edPR model, the S1 scenario is the same as that simulated for only a brief period in Fig 3. There, we observed that the ion concentrations gradually changed during the first seconds after the onset of stimulus (Fig 3E-H). However, for endured firing, the ion concentrations and reversal potentials settled on a (new) dynamic steady state (Fig 6E-J), where they deviated by ~ 1 mM from the baseline concentrations during rest (i.e., for edPR receiving no input). The apparent ‘thickness” of the curves (e.g., the orange curve for K^+^ in Fig 6I) is due to concentration fluctuations at the short time scale of AP firing. However, after each AP, the homeostatic mechanisms managed to re-establish ionic gradients before the next AP occurred, so that no slow concentration-dependent effect developed in the edPR model at a long time scale.

To simulate scenario S2, the PR model (Fig 7A-B) and edPR model (Fig 7C-J) were given a constant input (see figure caption) that gave them a firing rate of about 3 Hz. The PR model, which included no concentration-dependent effects, settled on a regular firing rate that it could maintain for an arbitrarily long time. Unlike the PR model, the edPR model did not settle at a steady state, but had a firing rate of ~ 3 Hz only for a period of ~ 5 s after stimulus onset. During this period, the ion concentrations gradually diverged from the baseline levels (Fig 7G-J). The corresponding changes in ionic reversal potentials (Fig 7E-F) affected the neuron’s firing properties and caused its firing rate to gradually increase before it eventually entered the depolarization block and got stuck at about *ϕ*_m_ =-30 mV. The main explanation behind the altered firing pattern was the change in the K^+^ reversal potential, which, for example, at 7 s after stimulus onset (*t* = 17 s) had increased by as much as 15 mV from baseline. This shift led to a depolarization of the neuron, which explains both the (gradually) increased firing rate and the (final) depolarization block, i.e., the condition where *ϕ*_m_ could no longer repolarize to levels below the firing threshold, and AP firing was abolished due to a permanent inactivation of active Na^+^ channels. Neuronal depolarization block is a well-studied phenomenon, which is often caused by high extracellular K^+^ concentrations [66].

**Figure 7.**
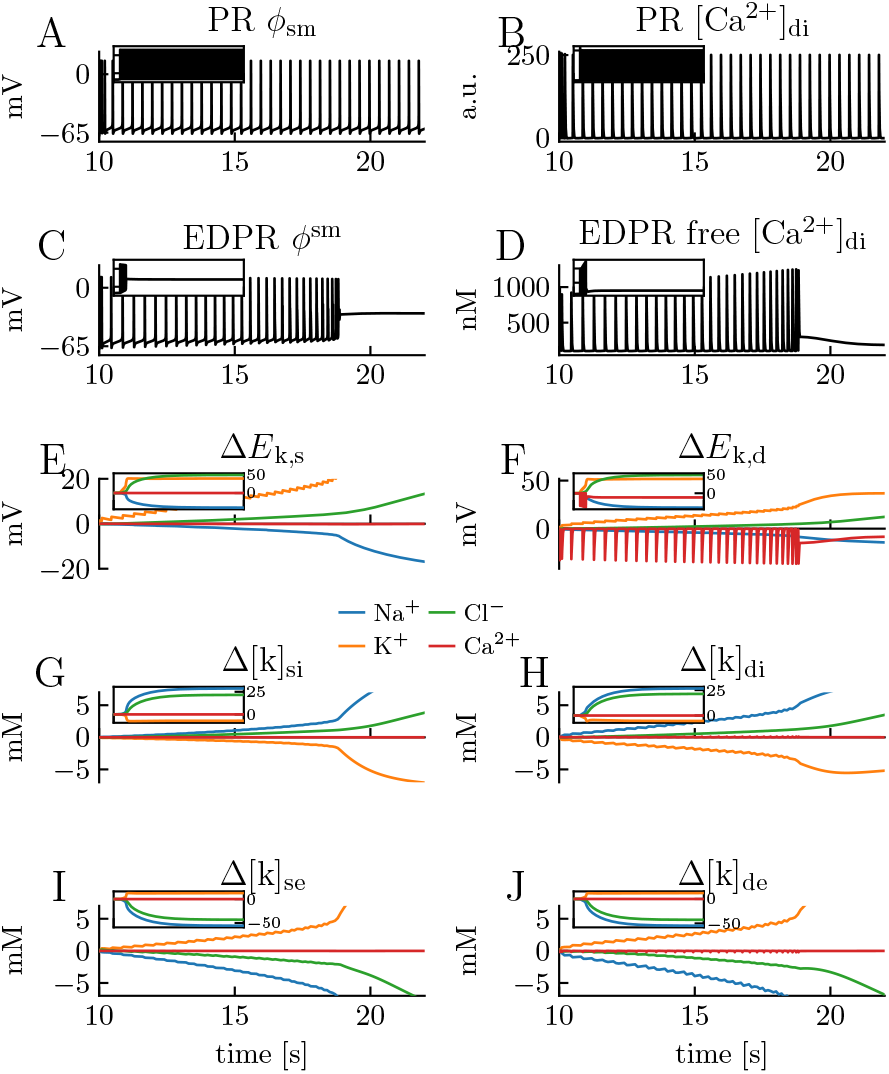
Model comparison for scenario with high frequency firing. Simulations on the PR model and edPR model when both models are driven by a constant input, giving them a firing rate of about 3 Hz. Simulations covered 200 s of biological time. **(A-D)** A 12 s sample of the dynamics of the somatic membrane potential *ϕ*_m_ and dendritic (free) Ca^2+^ concentration in the PR model **(A-B)** and edPR model **(C-D)**. The regular firing pattern in the PR model **(A-B)** was sustained over the full 200 s simulation (inset panels), while the edPR model stopped firing and entered depolarization block around *t* = 19 s. **(D)** Of the total amount of intracellular Ca^2+^, only 1% (as plotted) was assumed to be free (unbuffered). **(E-F)** Ionic reversal potentials and **(G-J)** ion concentrations in the edPR model varied throughout the simulation, and gradually diverged from baseline conditions. Indices *i, e, s*, and *d* indicate *intracellular, extracellular, soma*, and *dendrite*, respectively. Main panels show 12 s samples of the ion concentration dynamics, while insets show the dynamics over the full 200 s simulations. **(A-J)** Stimulus onset was *t* = 10 s in both models, and stimulus strength was *i*_stem_ = 1.55*μ*A/cm^2^ in the PR model **(A-B)** and *i*_stem_ = 46 pA in the edPR model **(C-J)**. See the Parameterizations section in Methods for a full description of the parameters used.

The homeostatic failure in S2 was due to the edPR model having a too high firing rate for the ion pumps and cotransporters to maintain ion concentrations close to baseline. The firing rate of 3 Hz was the limiting case (found by trial and error), i.e., for lower firing rates than this, the model could maintain regular firing for an arbitrarily long time. As many neurons can fire at quite high frequencies, a tolerance level of 3 Hz might seem a bit low, and we here provide some comments to this. Firstly, we note that the edPR model could fire at 3 Hz (and gradually higher frequencies) for about 9 s, and could also maintain a higher firing rate than this for a limited time. Secondly, the PR model, and thus the edPR model, represented a hippocampal CA3 neuron, which has been found to have an average firing rate of less than 0.5 Hz [67], so that endured firing of ≥ 3 Hz may be abnormal for these neurons. Thirdly, under biological conditions, glial cells, and in particular astrocytes, provide additional homeostatic functions [68] that were not accounted for in the edPR model, and the inclusion of such functions would probably increase the tolerance level of the neuron. Fourthly, the (3 Hz) tolerance level was a consequence of modeling choices and could be made higher, e.g., by increasing pump rates or compartment volumes. However, we did not do any model tuning in order to increase the tolerance level, as we, in light of the above arguments, considered a 3 Hz tolerance level to be acceptable.

### The edPR model predicts homeostatic failure due to impaired homeostatic mechanisms

Above we simulated homeostatic failure occurring because the firing rate became too high for the homeostatic mechanisms to keep up (S2). Homeostatic failure may also occur due to impairment of the homeostatic mechanisms, either due to genetic mutations (see, e.g., [69]) or because the energy supply is reduced, such as after a stroke (see, e.g., [25]). Here, we have used the edPR model to simulate an extreme version of this, i.e., a third scenario (S3) where all the homeostatic mechanisms were turned off.

In S3, the neuron received no external input, so that the dynamics of the neuron was solely due to gradually dissipating transmembrane ion concentration gradients. After an initial transient, we observed a slow and gradual increase in the membrane potential for about 30 s (Fig 8A). This coincided with a slow and gradual change in the ion concentrations (Fig 8D-G) and ionic reversal potentials (Fig 8B-C) due to predominantly passive leakage over the membrane.

**Figure 8.**
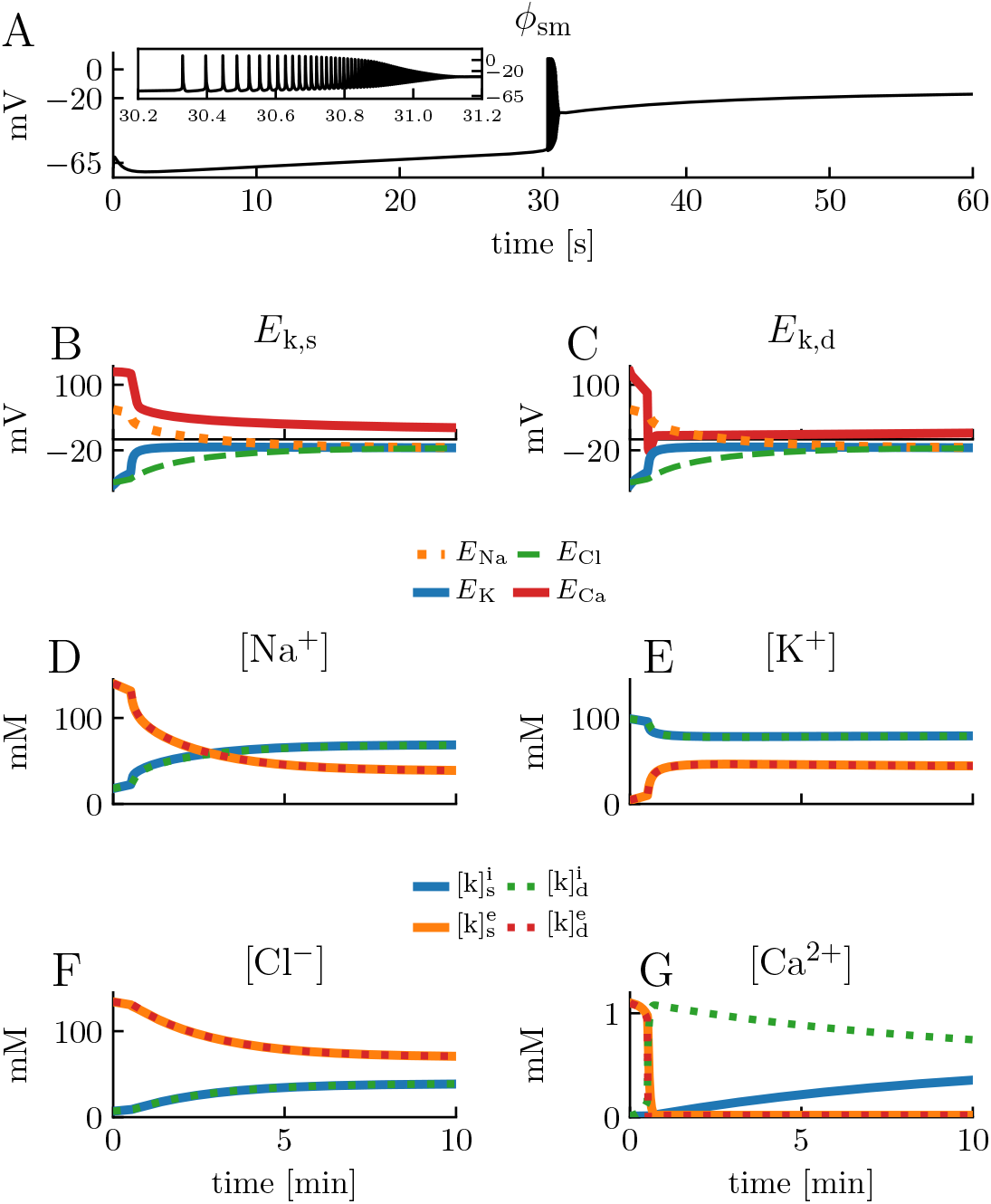
The wave of death. Simulations on the edPR model when all homeostatic mechanisms were turned off. The model received no external stimulus. Simulations covered 10 minutes of biological time. **(A)** A 60 s sample of the dynamics of the somatic membrane potential *ϕ*_m_. Inset shows a close-up of the burst of activity occurring at about *t* = 30 s. **(B-C)** Reversal potentials in the soma **(B)** and dendrite **(C)**. **(D-G)** Ion concentrations in all four compartments. Somatic and dendritic concentrations were almost identical for all ion species except for Ca^2+^. Indices *i, e, s*, and *d* indicate *intracellular, extracellular, soma*, and *dendrite*, respectively. See the Parameterizations section in Methods for a full description of the parameters used.

At about *t* = 30 s, the membrane potential reached the firing threshold, at which point the active channels started to use what was left of the concentration gradients to generate action potentials and Ca^2+^ spikes. This resulted in a burst of activity. During this bursts of activity, ion concentrations changed even faster, since both active and passive channels were then open. As a consequence, the “resting”membrane potential was further depolarized and the neuron went into depolarization block [66]. After this, the neuron continued to “leak” until it settled at a new steady state. The non-zero final equilibrium potential is known as the Donnan equilibrium or the Gibbs-Donnan equilibrium [70]. The reason why the cell did not approach an equilibrium with *ϕ*_m_ = 0 and identical ion concentrations on both side of the membrane, is that the model contained static residual charges, representing negatively charged macromolecules typically residing in the intracellular environment (see Methods), the sum of which resulted in a final state with a negatively charged inside. In addition, the membrane was also impermeable to Ca^2+^ in its final state (t > 1 min) since the Ca^2+^ channel inactivated, and the model contained no passive Ca^2+^ leakage. This explains why the Ca^2+^ reversal potential did not end up at Donnan-equilibrium potential, as did the reversal potential of the other mobile ions.

A pattern resembling that in Fig 8A, i.e., period of silence, followed by a burst of activity, and then silence again, has been seen in experimental EEG recordings of decapitated rats [71], where the activity burst was referred to as “the wave of death”, and the phenomenon was ascribed to the lack of energy supply to homeostatic mechanisms. The simulations in Fig 8A represents the single-cell correspondence to this death wave. We note that this phenomenon has been simulated and analyzed thoroughly in a previous modeling study, using a simpler, single compartmental model with ion conservation [40]. We, therefore, do not analyze it further here.

### Loss in accuracy when neglecting electrodiffusive effects on concentration dynamics

The concentration-dependent effects studied in the previous subsection were predominantly due to changes in ionic reversal potentials. Effects like this could therefore be accounted for by any model that in some way incorporates ion concentration dynamics [27–29, 33–57], provided that the ion concentration dynamics is accurately modeled. As we argued in the Introduction, previous multicompartmental neuron models that do incorporate ion concentration dynamics have not done it in a complete, ion conserving way that ensures a biophysically consistent relationship between ion concentration, electrical charge, and electrical potentials (see, e.g., [27, 48–57]). To specify, the change in the ion concentration in a given compartment will, in reality, depend on (i) the transmembrane influx of ions into this compartment, (ii) the diffusion of ions between this compartments and its neighboring compartment(s), and (iii) the electrical drift of ions between this compartment and its neighboring compartment(s). Some of the cited models account for only (i) [27, 49, 51], others account for (i) and (ii) [48, 50, 52–57], but neither account for (iii). When (iii) is not accounted for, electrical and diffusive processes are implicitly treated as independent processes, a simplifying assumption which is also incorporated in the reaction-diffusion module [72] in the NEURON simulation environment [73]. In models that apply this assumption, there will therefore be drift currents (along axons and dendrites) that affect *ϕ*_m_ (through the cable equation), but not the ion concentration dynamics, although they should, since also the drift currents are mediated by ions.

Here, we use simulations on the edPR model to test the inaccuracy introduced when not accounting for the effect of drift currents on ion concentration dynamics. We do so by comparing how many ions that were transferred from the somatic to the dendritic compartment through the intracellular (Fig 9A) and extracellular (Fig 9B) space, due to ionic diffusion (orange curves) versus electrical drift (blue curves), throughout the simulation in Fig 3. We note that Fig 9 shows the accumulatively moved number of ions (from time zero to *t*) due to axial fluxes exclusively. That is, the large number of, for example, Na^+^ ions transported intracellularly from the dendrite to the soma (negative sign) in Fig 9A1, does not by necessity mean that Na^+^ ions were piling up in the soma compartment, as the membrane efflux of Na^+^ was not accounted for in the figure.

**Figure 9.**
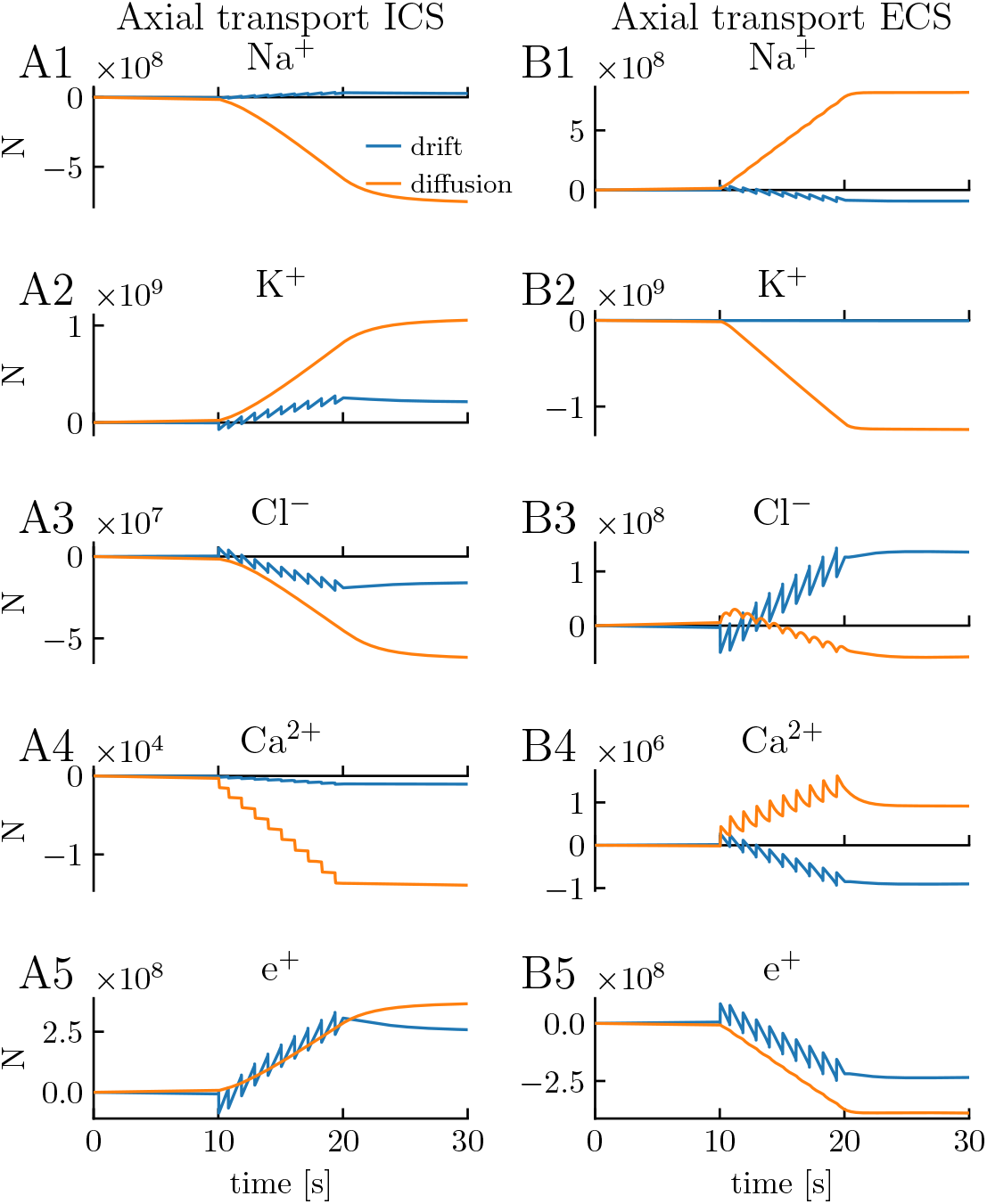
Axial transport of ions and charge due to drift versus diffusion. **(A1-A4)** The number of ions transported intracellularly from soma to dendrite from time zero to t by electrical drift versus ionic diffusion. **(B1-B4)** The number of ions transported extracellularly from (outside) soma to (outside) dendrite from time zero to t. **(A5)** Net charge transported intracellularly from soma to dendrite, represented as the number of unit charges e^+^. **(B5)** Net charge transported extracellularly from soma to dendrite, represented as the number of unit charges e^+^. **(A-B)** The simulation was the same as in Fig 3. See the Analysis section in Methods for a description of how we did the calculations.

Although diffusion tended to dominate the intracellular transport of ions on the long time scale (Fig 9A1-A4), the transport due to electrical drift was not vanishingly small. For example, the number of K^+^ and Cl^−^ ions transported by electrical drift was at the end of the stimulus period (*t* = 20 s) about 30 and 42 %, respectively, of the transport due to diffusion. In the extracellular space, diffusion was the clearly dominant transporter of Na^+^ and K^+^ (Fig 9B1-B2), while diffusion and electrical drift were of comparable magnitude for the other ion species (Fig 9B3-A4). Of course, these estimates are all specific to the ICRP model, as they will depend strongly on the included ion channels, ion pumps and cotransporters, and on how they are distributed between the soma and dendrite. In general, however, the simulations in Fig 9 suggest that electrical drift is likely to have a non-negligible effect on ion concentration dynamics, and that ignoring this effect will give rise to rather inaccurate estimates.

Finally, we also converted the sum of ionic fluxes in Fig 9 into an effective current, represented as the number of transported unit charges, e^+^ (Fig 9A5-B5). Interestingly, diffusion and drift contributed almost equally to the axial charge transport in the system. We note, however, that the movement of charges per time unit is indicated by the slope of the curves, which was much larger for the drift case (blue curve) than for diffusion (orange curve). The drift curve had a jigsaw shape, which shows that drift was moving charges back and forth in the system, while the diffusion always went in the same direction, explaining why it, despite being smaller than the drift current, had a comparably large accumulative effect on charge transport. The temporally averaged picture of charge transport that emerges from Fig 9A5 is that of a slow current loop where charge is transferred intracellularly from the soma to the dendrite (Fig 9A5), where it crosses the membrane (outward current), and then is transferred extracellularly back from the dendrite to the soma (Fig 9B5), before crossing the membrane again (inward current). This configuration is similar to the slow loop current seen during spatial buffering by astrocytes (see, e.g. Fig 1 in [68]).

### Loss in accuracy when neglecting electrodiffusive effects on voltage dynamics

In the previous section, we investigated the consequences of neglecting (iii) the contribution of drift currents on ion concentration dynamics. Here, we investigate the consequences of neglecting the effect of ionic diffusion (along dendrites and axons) on the electrical potential, focusing on the extracellular potential *ϕ*_e_. Forward modeling of extracellular potentials is typically based on volume conductor (VC) theory [16–18, 20, 21], which assumes that diffusive effects on electrical potentials are negligible. Being based on a unified electrodiffusive KNP framework (Fig 1), the edPR model accounts for the effects of ionic diffusion on the electrical potentials, and can thus be used to address the validity of this assumption.

To illustrate the effect of diffusion on *ϕ*_e_, we may split it into a component *ϕ*_VC,e_ explained by standard VC-theory, and a component *ϕ*_diff,e_ representing the additional contribution caused by diffusive currents (Eq 81). In the simulation in Fig 3, the diffusive contribution was found to be very small compared to the VC-component (Fig 10). However, while *ϕ*_VC,e_ fluctuated rapidly from negative to positive values during neuronal activity, *ϕ*_diff,e_ varied on a slower time scale and had the same directionality throughout the simulation. This is equivalent to what we saw in Fig 9B5, i.e., that diffusion always moved charge in the same direction. As we also have shown in previous studies, diffusion is thus likely to be important for the slow (direct-current (DC) like) effects on extracellular potentials [31, 32, 74, 75]. Albeit small, the slowly varying diffusion evoked shifts in *ϕ*_e_ are putatively important for explaining the DC-shifts and long-time concentration dynamics reported during, e.g., spreading depression [25, 26].

**Figure 10.**
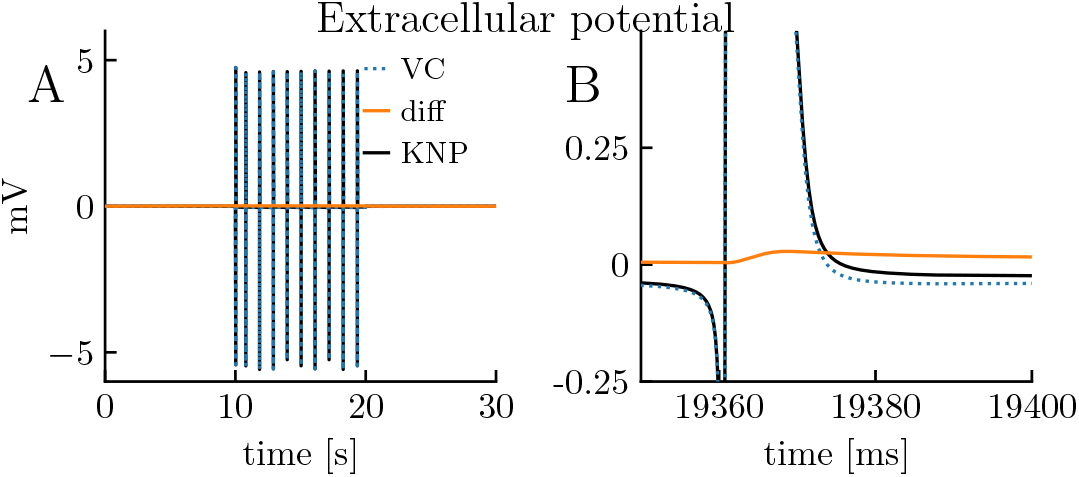
Effect of diffusion on extracellular potential. The extracellular potential *ϕ*_e_ in the edPR model, split (cf. Eq 81) into a component explained by standard VC-theory (*ϕ*_VC,e_) and a ‘‘correction” (*ϕ*_diff,e_) when diffusive contributions are accounted for. **(A-B)** The simulation was the same as in Fig 3. **(B)** Close-up of selected AP in **(A)**. See the Analysis section in Methods for a description of how we calculated *ϕ*_VC,e_ and *ϕ*_diff,e_. mean(*ϕ*_e_)=−0.0021 mV, mean(*ϕ*_diff,e_) = 0.0034 mV, mean(*ϕ*_VC,e_)=−0.0055 mV

## Discussion

The original Pinsky-Rinzel (PR) is a reduced model of a hippocampal neuron, which reproduces the essential somatodendritic firing properties of CA3 neurons despite having only two compartments [3]. Simplified neuron models like that are useful, partly because their reduced complexity makes them easier to analyze, and as such, can lead to insight in key neuronal mechanisms, and partly because they demand less computer power and can be used as modules in large scale network simulations. Whereas the PR model, as most available neuron models, assumes that ion concentrations remain constant during the simulated period, the electrodiffusive Pinsky-Rinzel (edPR) proposed here models ion concentration dynamics explicitly. The edPR model may thus be seen as a supplement to the PR model, which should be applied to simulate conditions where ion concentrations are expected to vary with time.

In the results section, we showed that the edPR model closely reproduced the firing properties of the PR model for short term dynamics (Fig 5), and for long term dynamics provided that the firing rate was sufficiently low for the homeostatic mechanisms to maintain ion concentrations close to baseline (Fig 6). We also showed that if the firing rate became too high (Fig 7), or if the homeostatic mechanisms were impaired (Fig 8), unsuccessful homeostasis would cause ion concentrations to gradually shift over time, and lead to slowly developing changes in the firing properties of the edPR model, changes that were not accounted for by the original PR model. The edPR model was based on an electrodiffusive framework [60], which ensured a consistent relationship between ion concentrations, electrical charge, and electrical potential in four compartments. To our knowledge, the edPR model is the first multicompartmental neuronal model that ensures a complete and consistent ion concentration and charge conservation.

### Model assumptions

The construction of the edPR model naturally involved making a set of modeling choices, and the most important of these are discussed here. Firstly, in the construction of the model, we focused on morphological simplicity, biophysical rigor, and mechanistic understanding, rather than on replicating any specific biological scenario and incorporating biological details. Secondly, simultaneous data of variations in all intra- and extracellular concentrations during neuronal firing are not available, and it might not even be feasible to obtain such data. Consequently, modeling based on biophysical constraints may be the best means to estimate it. The concentration dynamics in the edPR model were thus not directly constrained to data but constrained so that there was, at all times, an internally consistent relationship between all ion concentrations and all electrical potentials, ensuring an electroneutral bulk solution. Thirdly, to include extracellular dynamics to models of neurons or networks of such is computationally challenging, since the extracellular space, in reality, is an un-confined three-dimensional continuum, locally affected by populations of nearby neurons and glial cells. As we wanted to keep things simple and conceptual, we chose to use closed boundary conditions, i.e., no ions and no charge were allowed to leave or enter the system consisting of the single (2-compartment) neuron and its local and confined (2-compartment) surrounding (Fig 2).

A consequence of using closed boundary conditions was that the extracellular (like the intracellular) currents became one-dimensional (from soma to dendrite), while in reality, extracellular currents pass through a three-dimensional volume conductor. The edPR model could be made three dimensional if embedded in a bi- or tri-domain model (as discussed below). However, currently, it is 1D, and the effect of the 1D assumption was essentially an increase in the total resistance (fewer degrees of freedom) for extracellular currents, which gave rise to an artificially high amplitude in extracellular AP signatures (Fig 3). We note, however, that the closed boundary is actually equivalent to assuming periodic boundary conditions, so that the edPR model essentially simulates the hypothetical case of a population of perfectly synchronized neurons, i.e., one where all neurons are doing exactly the same as the simulated neuron, so that no spatial variation occurs. Likely, this may give accurate predictions for ion concentration shifts over time, as these reflect a temporal average of activity, but less accurate predictions for brief and unique electrical events, such as action potentials, which are not likely to be elicited in perfect synchrony by large population [31].

Fourthly, to faithfully represent a morphologically complex neuron with a reduced number of compartments is a non-trivial task. Available analytical theory for collapsing branching dendrites into equivalent cylinders are generally based on certain assumptions about branching symmetries, and on preserving electrotonic distances [76]. However, it is unlikely that the length constants of electrodynamics and ion concentration dynamics scale in the same way. Hence, in the edPR model, the volumes and membrane areas of-, and cross-section areas between, the two neuronal compartments were here introduces as rather arbitrary model choices, fixed at values that were verified to give agreement between the firing properties of the edPR model and the PR model.

### Outlook

Being applicable to simulate conditions with failed homeostasis, the edPR model opens up for simulating a range of pathological conditions, such as spreading depression or epilepsy [22–25], which are associated with large scale shifts in extracellular ion concentrations. A particular context in which we anticipate the edPR model to be useful is that of simulating spreading depression. Previous spatial, electrodiffusive, and biophysically consistent models of spreading depression have targeted the problem at a large-scale tissue-level, using a mean-field approach [30, 77, 78]. These models were inspired by the *bi-domain* model [79], which has been successfully applied in simulations of cardiac tissue [80, 81]. The bi-domain model is a coarse-grained model, in which the tissue is considered as a bi-phasic continuum consisting of an intracellular and extracellular domain. That is, a set of intra- and extracellular variables (i.e., voltages and ion concentrations), and the ionic exchange between the intra- and extracellular domains, are defined at each point in space. This simplification allows for large scale simulations of signals that propagate through tissue but sacrifices morphological detail. In the context of spreading depression, a shortcoming with this simplification is that the leading edge of the spreading depression wave in both the hippocampus and cortex is in the layers containing the apical dendrites [22]. This suggests that the different expression of membrane mechanisms in deeper (somatic) and higher (dendritic) layers may be crucial for fully understanding the propagation and genesis of the wave. In this context, the edPR model could enter as a module in a, let us say, *bi-times-two-domain* model, where each point in (*xy*) space contains a set of (i) somatic intracellular variables, (ii) somatic extracellular variables, (iii) dendritic intracellular variables and (iv) dendritic extracellular variables, and thus accounts for the differences between the higher and lower layers. We should note that the state of the art models of spreading depression are not bi-domain models but rather tri-domain models, as they also include a glial domain to account especially for the work done by astrocytes in K^+^ buffering [30, 77, 78]. Hence, to use the edPR model to expand the current spreading depression models, a natural first step would be to include a glial (astrocytic) compartment in it, so that it eventually could be implemented as a *tri-times-two-domain* model.

## Methods

### The Kirchoff-Nernst-Planck (KNP) framework

In the following section, we derive the KNP continuity equations for a one-dimensional system containing two plus two compartments (Fig 2A), with sealed boundary conditions (i.e., no ions can enter or leave the system). The geometrical parameters used in the edPR model were as defined in Table 1.

**Table 1.** Geometrical parameters

| Parameter | Value |
| --- | --- |
| $\Delta x$ (distance between the two compartments) | $667 \cdot 10^{-6} \text{ m}$ |
| $A_s$ (somatic membrane area) | $616 \cdot 10^{-12} \text{ m}^2$ |
| $A_d$ (dendritic membrane area) | $616 \cdot 10^{-12} \text{ m}^2$ |
| $A_i$ (intracellular cross-section area) | $\alpha \cdot A_s^\dagger$ |
| $A_e$ (extracellular cross-section area) | $A_i/2$ |
| $V_{si}$ (somatic intracellular volume) | $1437 \cdot 10^{-18} \text{ m}^3$ |
| $V_{se}$ (somatic extracellular volume) | $1437 \cdot 10^{-18} \text{ m}^3$ |
| $V_{di}$ (dendritic intracellular volume) | $718.5 \cdot 10^{-18} \text{ m}^3$ |
| $V_{de}$ (dendritic extracellular volume) | $718.5 \cdot 10^{-18} \text{ m}^3$ |

The intracellular volumes ( $V_{si}$ , $V_{di}$ ) and membrane areas ( $A_s$ , $A_d$ ) correspond to spheres with radius $7 \mu\text{m}$ . We used the same intra-/extracellular volume ratio as in [40].
<sup>†</sup> The parameter $\alpha$ describes the coupling strength of the model and is defined in the Parameterizations section. Its default value was 2.

Two kinds of fluxes are involved: transmembrane fluxes and intra- and extracellular fluxes. The framework is general to the choice of the transmembrane fluxes. A transmembrane flux of ion species k (*j*_k,m_) represents the sum of all fluxes through all membrane mechanisms that allow ion k to cross the membrane.

Intracellular flux densities are described by the Nernst-Planck equation:

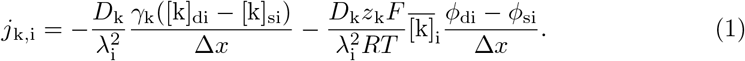

In Eq 1, *D*_k_ is the diffusion constant, γ_k_ is the fraction of freely moving ions, that is, ions that are not buffered or taken up by the ER, λ_i_ is the tortuosity, which represents the slowing down of diffusion due to obstacles, γ_k_([k]_di_ − [k]_si_)/Δ*x* is the axial concentration gradient, *z*_k_ is the charge number of ion species k, *F* is the Faraday constant, *R* is the gas constant, *T* is the absolute temperature, 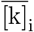 is the average concentration, that is, γ_k_ ([k]_di_ + [k]_si_)/2, and (*ϕ*_di_ − *ϕ*_si_)/Δ*x* is the axial potential gradient. Similarly, the extracellular flux densities are described by

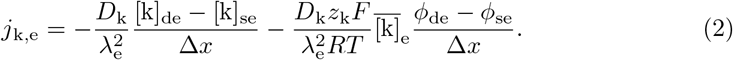

In Eq 2, we do not include γ_k_, as all ions can move freely in the extracellular space. Diffusion constants, tortuosities, and intracellular fractions of freely moving ions used in the edPR model were as in Table 2.

**Table 2.** Diffusion constants, tortuosities, and intracellular fractions of freely moving ions

| Parameter | Value | Reference |
| --- | --- | --- |
| $D_{\text{Na}}$ ($\text{Na}^+$ diffusion constant) | $1.33 \cdot 10^{-9} \text{ m}^2/\text{s}$ | [31, 82] |
| $D_{\text{K}}$ ($\text{K}^+$ diffusion constant) | $1.96 \cdot 10^{-9} \text{ m}^2/\text{s}$ | [31, 82] |
| $D_{\text{Cl}}$ ($\text{Cl}^-$ diffusion constant) | $2.03 \cdot 10^{-9} \text{ m}^2/\text{s}$ | [31, 82] |
| $D_{\text{Ca}}$ ($\text{Ca}^{2+}$ diffusion constant) | $0.71 \cdot 10^{-9} \text{ m}^2/\text{s}$ | [31, 82] |
| $\lambda_i$ (intracellular tortuosity) | 3.2 | [60, 83] |
| $\lambda_e$ (extracellular tortuosity) | 1.6 | [60, 83] |
| $\gamma_{\text{Na}}, \gamma_{\text{K}}, \gamma_{\text{Cl}}$ (intracellular fractions of free ions) | 1 | |
| $\gamma_{\text{Ca}}$ (intracellular fraction of free ions) | 0.01 | |

### Ion conservation

The KNP framework is based on the constraint of ion conservation. To keep track of ion concentrations we solve four differential equations for each ion species k:

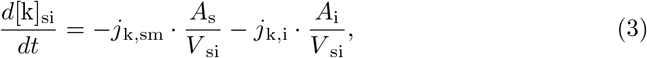

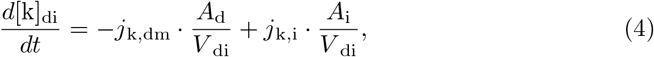

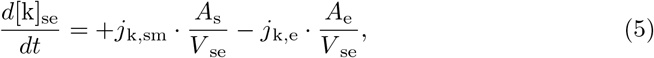

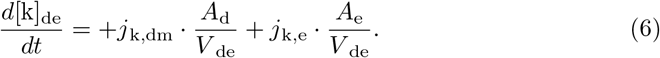

For each compartment, all flux densities are multiplied by the area they go through and divided by the volume they enter to calculate the change in ion concentration. If we insert the Nernst-Planck equation (Eq 1) for the intracellular flux density, the first of these equations becomes:

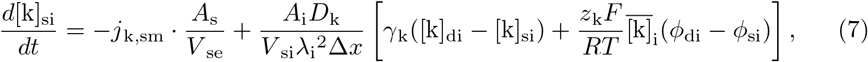

where the voltage variables so far are undefined.

### Four constraints to derive *ϕ*

If we have four ion species (Na^+^, K^+^, Cl^−^, and Ca^2+^) in four compartments, we have 20 unknown parameters (16 for [k] and four for *ϕ*), while Eqs 3-6 for four ion species give us only 16 equations. To solve this, we need to define additional constraints that allow us to express the potentials *ϕ* in terms of ion concentrations.

#### Arbitrary reference point for *ϕ*

As we may define an arbitrary reference point for *ϕ*, we take

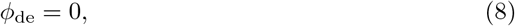

as our first constraint, i.e., the potential outside the dendrite is defined to be zero.

#### (ii) Membrane is a parallel plate capacitor

The second constraint is that the membrane is a parallel plate capacitor that always separates a charge *Q* on one side from an opposite charge −*Q* on the other side, giving rise to a voltage difference

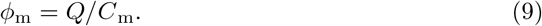

Here, *C*_m_ is the total capacitance of the membrane, i.e., *C*_m_ = *c*_m_*A*_m_, where *c*_m_ is the more commonly used capacitance per membrane area. As, by definition, *ϕ*_m_ = *ϕ*_i_ − *ϕ*_e_, we get:

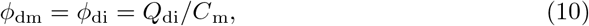

in the dendrite (since *ϕ*_de_ = 0), and

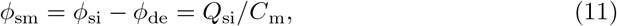

in the soma.

#### (iii) Bulk electroneutrality

The KNP scheme is based on the assumption of bulk electroneutrality, which means the net charge associated with the ion concentrations in a given compartment by constraint must be identical to the membrane charge in this compartment. The intracellular dendritic charge is thus 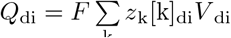. By inserting this into Eq 10, we obtain the final expression for *ϕ*_di_:

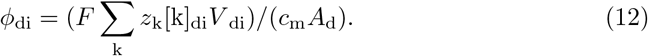

By inserting the equivalent expression for *Q*_si_ into Eq 11, we get

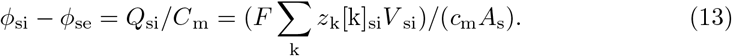

Here, the extracellular potential is not set to zero, so we need a fourth constraint to determine *ϕ*_si_ and *ϕ*_se_ separately.

#### (iv) Current anti-symmetry

For the charge anti-symmetry between the two sides of the capacitive membrane (*Q*_i_ = −*Q*_e_) to be preserved in time, we must define our initial conditions so that this is the case at *t* = 0, and the system dynamics so that this stays the case. Hence, the system dynamics must ensure that *dQ*_di_/*dt* = −*dQ*_de_/*dt* and *dQ*_si_/*dt* = −*dQ*_se_/*dt*. The membrane currents (in isolation) will always fulfill this criterion, as any charge that crosses the membrane by definition disappears from one side of it and pops up at the other. Hence, we thus need to make sure that also the axial currents (in isolation) fulfill the criterion. The system must thus be constrained so that, if an extracellular current transports a charge *δq* into a given extracellular compartment, the intracellular current must transport the opposite charge −*δq* into the adjoint intracellular compartment. That is, we must have that:

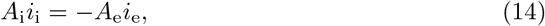

where *i*_1_ and *i*_e_ are the intra- and extracellular current densities, respectively. To find an expression for these, we multiply Eqs 1 and 2 by *Fz*_k_ and sum over all ion species k. The expressions for the intra- and extracellular current densities then become:

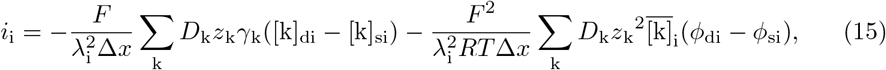

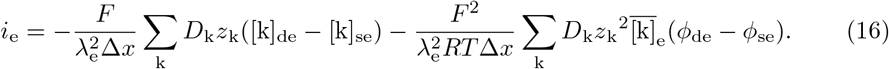

In Eq 15, the first term is the diffusion current density:

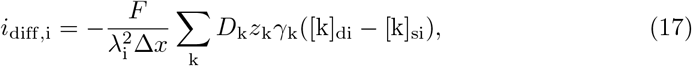

which is defined by the ion concentrations. The second term is the field driven current density

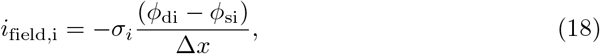

where we have identified the conductivity as

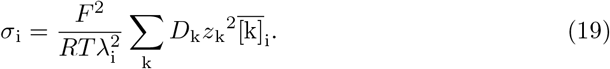

Similarly, Eq 16 can be written in terms of *i*_diff,e_, *i*_fieid,e_, and *σ*_e_. By combining Eqs 14, 15, and 16, we obtain:

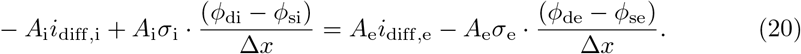

In Eq 20, *ϕ*_di_ and *ϕ*_de_ are already known from Eqs 8 and 12, while *i*_diff_ and *σ* are expressed in terms of ion concentrations. We may thus solve Eqs 13 and 20 for the last two voltage variables *ϕ*_se_ and *ϕ*_si_:

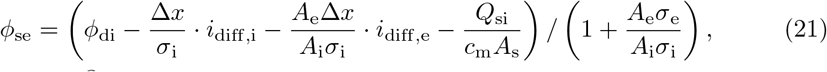

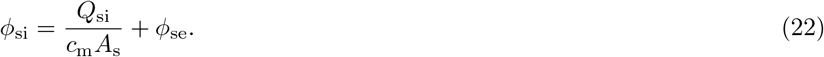

### Membrane mechanics

#### Leakage channels

In the original PR model, the membrane leak current represents the combined contribution from all ion species. When using the KNP framework, on the other hand, where we keep track of all ions separately, the leak current must be ion-specific. We modeled this as in [45], that is, for each ion species k, we implemented a passive flux density across the membrane

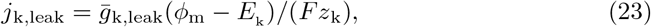

where *ḡ*_k,leak_ is the conductance, *ϕ*_m_ is the membrane potential, *E*_k_ is the reversal potential, *F* is the Faraday constant, and *z*_k_ is the charge number. The reversal potential is a function of ion concentrations, and is calculated using the Nernst equation:

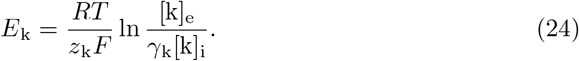

Here, *R* is the gas constant, *T* is the absolute temperature, γ_k_ is the intracellular fraction of free ions, and [k]_e_ and [k]_i_ are the concentrations of ion k outside and inside the cell, respectively. We included Na^+^, K^+^, and Cl^−^ leak currents in both compartments.

#### Active ion channels

All active ion channel currents were adopted from the original PR model [3], as they were described in [8], and converted to ion channel fluxes. The soma compartment contained a Na^+^ flux (*j*_Na_) and a K^+^ delayed rectifier flux (*j*_K–DR_), while the dendrite contained a voltage-dependent Ca^2+^ flux (*j*_Ca_), a voltage-dependent K^+^ AHP flux (*j*_K–AHP_), and a Ca^2+^-dependent K^+^ flux (*j*_K–C_):

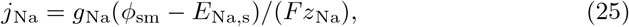

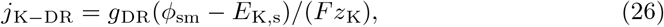

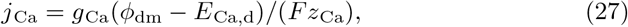

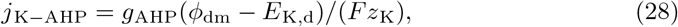

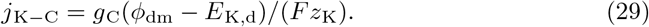

The voltage-dependent conductances were modeled using the Hodkin-Huxley formalism with differential equations for the gating variables:

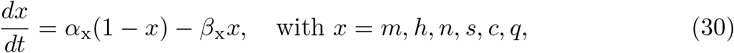

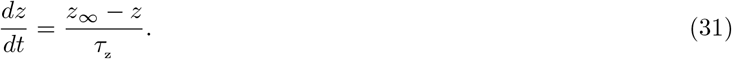

The conductances and gating variables were given by:

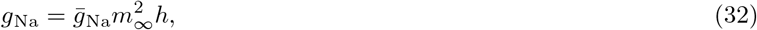

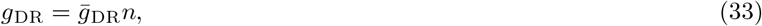

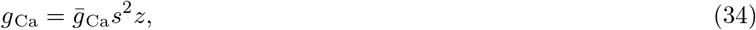

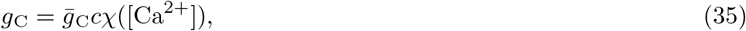

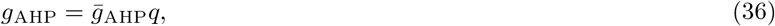

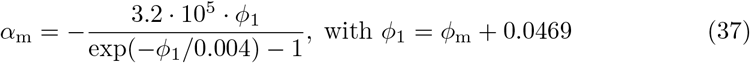

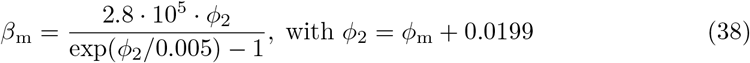

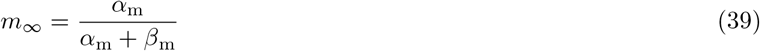

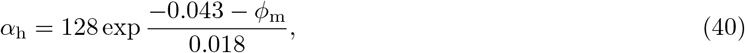

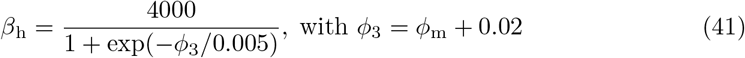

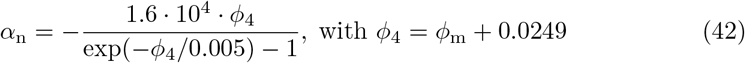

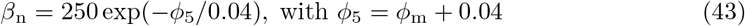

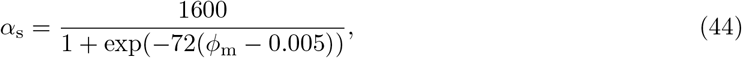

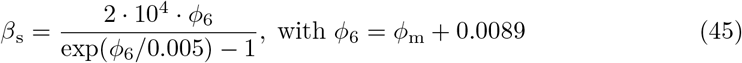

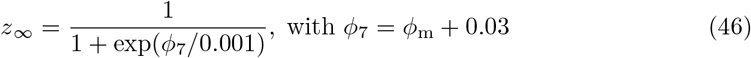

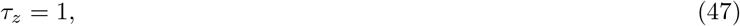

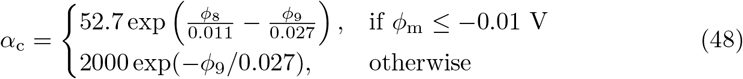

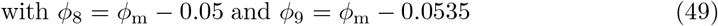

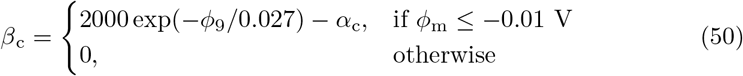

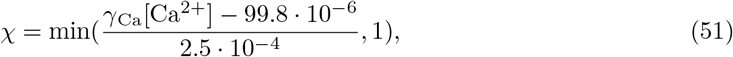

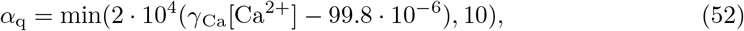

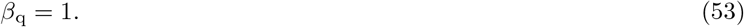

All these equations were taken from [8] (with errata [84]), and converted so that values are given in SI units: units for rates (*α*’s, *β*’s, *χ*) are 1/s, units for *τ_z_* is s, and voltages *ϕ* should be inserted with units in V. The equations were used in their original form, except those related to Ca^2+^ dynamics, where we made the following changes: Firstly, as a large fraction of intracellular Ca^2+^ is buffered or taken up by the ER, we multiplied [Ca^2+^] in Eqs 51 and 52 by a factor γ_Ca_, which refers to the fraction of free Ca^2+^ within the cell, and set this to be 0.01. As [Ca^2+^] in Eqs 51 and 52 were multiplied with 0.01 only the free Ca^2+^ could affect the Ca^2+^ activated ion channels. We further assumed that only the free Ca^2+^ could move between the intracellular compartments (Eq 1) and affect the Ca^2+^ reversal potential (Eq 24). Secondly, the original PR model had an abstract and unitless variable for the intracellular Ca^2+^ concentration, with a basal concentration of 0.2, while we defined a (biophysically realistic) baseline concentration of 0.01 mM, which corresponds to a concentration of *free* Ca^2+^ of 100 nM. In Eqs 51 and 52 we therefore subtracted 99.8 o 10^-6^(mol/m^3^) from the Ca^2+^ concentration to correct for the shift in baseline. Thirdly, we modified the voltage-dependent Ca^2+^ current to include an inactivation variable *z* (Eqs 31 and 34). We implemented this inactivation like they did in [85] (Eqs. A2-A3), but set the time constant *τ*_z_ to 1 s, the half-activation voltage to −30 mV, and the slope of the steady-state Boltzmann fit to *z*_∞_ to 0.001. In the original PR model, inactivation was neglected due to the argument that it was too slow to have an impact on simulation outcomes [2]. However, in our simulations, we observed that it had a significant impact, and therefore we included it.

#### Homeostatic mechanisms

To maintain baseline ion concentrations for low frequency activity we added a 3Na^+^/2K^+^ pump, a K^+^/Cl^−^ cotransporter (KCC2), and a Na^+^/K^+^/2Cl^−^ cotransporter (NKCC1). Their functional forms were taken from [45].

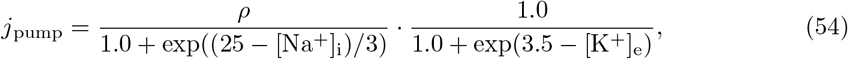

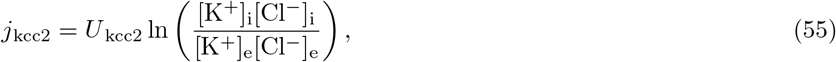

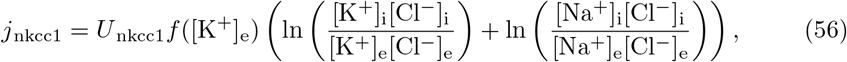

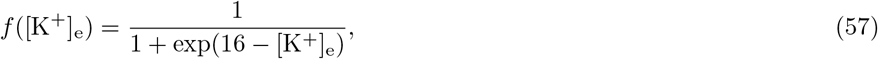

where *ρ*, *U*_kcc2_, and *U*_nkcc1_ are pump and cotransporter strengths. We assumed optimal pump functionality and set *ρ* to be the pump strength used in [45] for the fully oxygenated state with normal bath potassium (*ρ*_max_).

Intracellular Ca^2+^ decay was modeled in a similar fashion as in [3], but to ensure ion conservation we modeled it as Ca^2+^/Na^+^ exchanger, exchanging one Ca^2+^ (outward) for two Na^+^ (inward). The Ca^2+^ decay flux density was defined as:

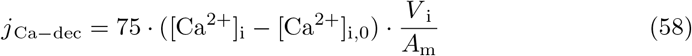

where 75 is the decay rate, same as in [3] but in SI units, and [Ca^2+^]_i,0_ is the initial Ca^2+^ concentration.

#### Model summary

We summarize the model here for easy reference. In short, we solved four differential equations for all ion species k:

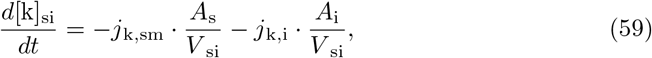

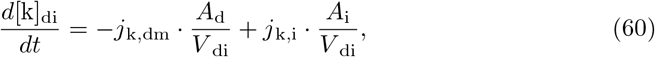

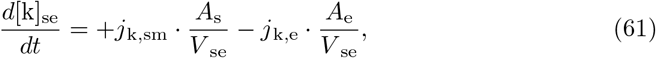

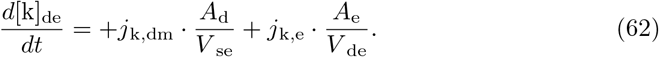

At each time step, *ϕ* in all four compartments was derived algebraically:

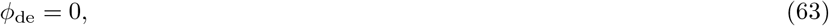

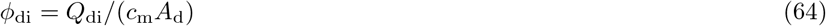

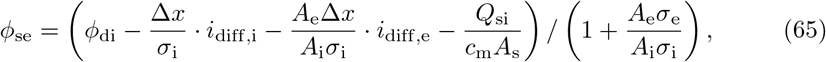

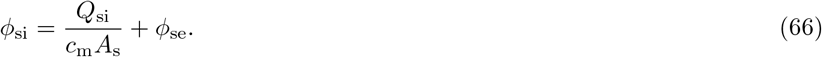

The total membrane flux densities were as follows:

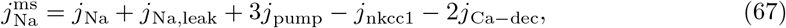

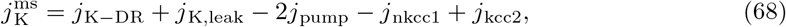

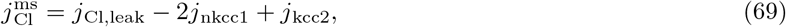

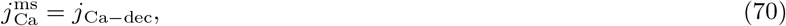

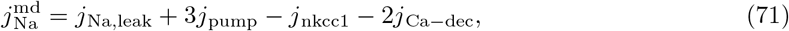

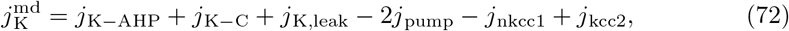

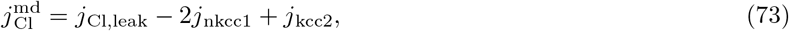

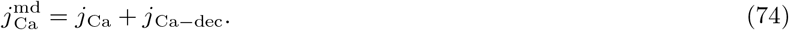

Figure 2 summarizes the model. The parameters involved in this model and their values used in this study are listed in Tables 1-4.

**Table 3.**
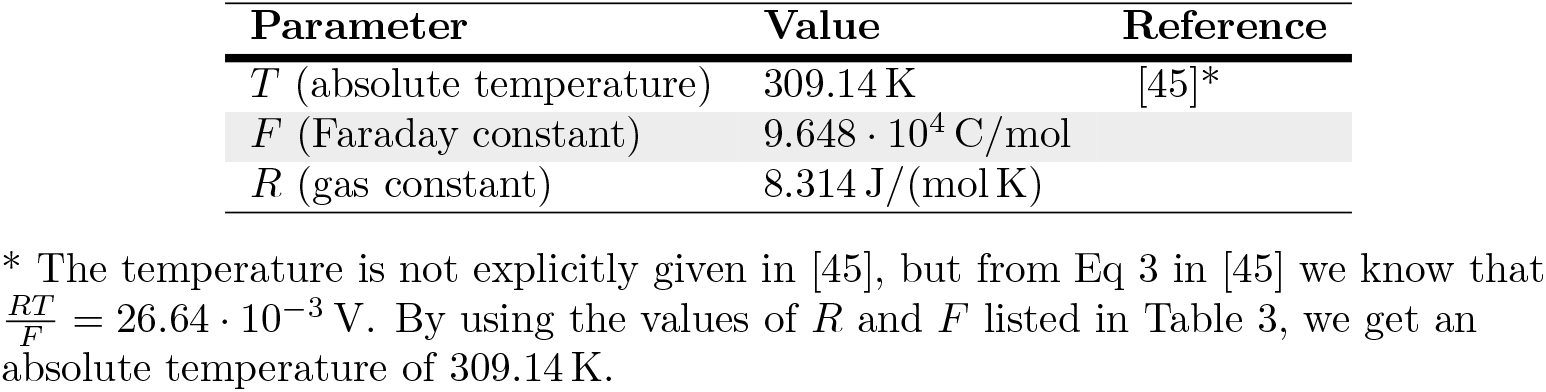
Temperature and physical constants

| Parameter | Value | Reference |
| --- | --- | --- |
| $T$ (absolute temperature) | 309.14 K | [45]* |
| $F$ (Faraday constant) | $9.648 \cdot 10^4$ C/mol | |
| $R$ (gas constant) | 8.314 J/(mol K) | |

\* The temperature is not explicitly given in [45], but from Eq 3 in [45] we know that $\frac{RT}{F} = 26.64 \cdot 10^{-3}$ V. By using the values of $R$ and $F$ listed in Table 3, we get an absolute temperature of 309.14 K.

### Original Pinsky-Rinzel model

We implemented the original Pinsky-Rinzel equations from Box 8.1 in [8]. The reversal potential of the leak current, not specified in [8], was set to —68 mV to ensure a resting potential close to that of the edPR model. We also used this as the initial potentials, that is, *ϕ*_sm,0_ = –68 mV and *ϕ* _dm,0_ = –68 mV. The other initial conditions were *n*_0_ = 0.001, *h*_0_ = 0.999, *s*_0_ = 0.009, *c*_0_ = 0.007, *q*_0_ = 0.01, and [Ca^2+^]_0_= 0.2, same as in [3].

### Simulations

#### Parameterizations

The parameters listed in Tables 1-4 were used in all the simulations of the electrodiffusive Pinsky-Rinzel (edPR) model. We tuned the Ca^2+^ conductance *ḡ*_Ca_ manually to obtain comparable spike shapes between the edPR model and the original PR model, as well as the fraction of free Ca^2+^ inside the cell, and the coupling strength between the soma and the dendrite.

**Table 4.** Membrane parameters

| Parameter | Value | Reference |
| --- | --- | --- |
| $c_m$ | $3 \cdot 10^{-2} \text{ F/m}^2$ | [3, 8] |
| $\bar{g}_{\text{Na,leak}}$ | $0.247 \text{ S/m}^2$ | [45] |
| $\bar{g}_{\text{K,leak}}$ | $0.5 \text{ S/m}^2$ | [45] |
| $\bar{g}_{\text{Cl,leak}}$ | $1.0 \text{ S/m}^2$ | [45] |
| $\bar{g}_{\text{Na}}$ | $300 \text{ S/m}^2$ | [3, 8] |
| $\bar{g}_{\text{DR}}$ | $150 \text{ S/m}^2$ | [3, 8] |
| $\bar{g}_{\text{Ca}}$ | $118 \text{ S/m}^2$ | |
| $\bar{g}_{\text{AHP}}$ | $8 \text{ S/m}^2$ | [3, 8] |
| $\bar{g}_{\text{C}}$ | $150 \text{ S/m}^2$ | [3, 8] |
| $\rho$ | $1.87 \cdot 10^{-6} \text{ mol}/(\text{m}^2\text{s})$ | [45]* |
| $U_{\text{kcc2}}$ | $7.0 \cdot 10^{-7} \text{ mol}/(\text{m}^2\text{s})$ | [45]* |
| $U_{\text{nkcc1}}$ | $2.33 \cdot 10^{-7} \text{ mol}/(\text{m}^2\text{s})$ | [45]* |

\* We multiplied the original values from [45] by a conversion factor $\frac{7}{3} \cdot 10^{-6} \text{ m}$ to convert the units from mM/s to mol/m<sup>2</sup>s. The conversion factor equals the initial inverse surface area to volume ratio from [45].

In the edPR model, the coupling strength between the soma and dendrite was proportional to the ratio *A*_i_/Δ*x*, and all model outputs depended on this ratio, and not on *A*_i_ or Δ*x* in isolation. By choice, we adjusted the coupling strength by varying *A*_i_ = *αA*_m_ through adjusting the parameter *α*. We could have obtained the equivalent effect by varying Δ*x* instead. The default value of *α* was set to 2. All simulations were run using this value, except in Fig 5C where *α* was set to 0.43.

In the original PR model, the coupling strength between the soma and dendrite was represented by a coupling conductance *g_c_*, which had a default value of 10.5mS/cm^2^. In Fig 5A, *g_c_* was set to 2.26mS/cm^2^.

#### Initial conditions

The initial conditions of the edPR model are listed in Table 5. We adjusted the initial ion concentrations manually to ensure a stable resting state of the model. Their values give us the following reversal potentials: *E*_Na_ = 55 mV, *E*_K_ = —84 mV, *E*_Cl_ = —79 mV, and *E*_Ca_ = 124 mV. The variables [k_res_]_i_ and [k_res_]_e_ are static residual charges. They represent negatively charged macromolecules typically residing in the intracellular environment. We defined them as

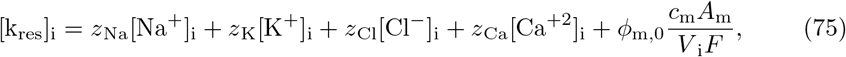

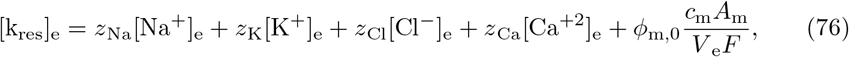

**Table 5.** Initial conditions

| Variables | Value |
| --- | --- |
| $[\text{Na}^+]_{\text{i}}$ | 18 mM |
| $[\text{Na}^+]_{\text{e}}$ | 140 mM |
| $[\text{K}^+]_{\text{i}}$ | 99 mM |
| $[\text{K}^+]_{\text{e}}$ | 4.3 mM |
| $[\text{Cl}^-]_{\text{i}}$ | 7 mM |
| $[\text{Cl}^-]_{\text{e}}$ | 134 mM |
| $[\text{Ca}^{+2}]_{\text{i}}$ | 0.01 mM* |
| $[\text{Ca}^{+2}]_{\text{e}}$ | 1.1 mM |
| $[k_{\text{res}}]_{\text{i}}$ | $-110$ mM<sup>†</sup> |
| $[k_{\text{res}}]_{\text{e}}$ | $-12$ mM<sup>†</sup> |
| $n$ | 0.0003 |
| $h$ | 0.999 |
| $s$ | 0.007 |
| $c$ | 0.006 |
| $q$ | 0.011 |
| $z$ | 1.0 |

\* Only 1% of this total intracellular $\text{Ca}^{2+}$ , that is, a 100 nM, was assumed to be free (unbuffered).
<sup>†</sup> $-110$ mM and $-12$ mM are approximate values. Exact values were calculated from Eqs. 75 and 76.

where *φ*_m,0_ is the initial membrane voltage, set to –68 mV in all simulations. The initial conditions were equal in both the somatic and the dendritic compartment.

#### Stimulus current

We stimulated the cell by injecting a K^+^ current *i*_stim_ into the soma. Previous computational modeling of a cardiac cell has shown that stimulus with K^+^ causes the least physiological disruption [33]. To ensure ion conservation, we removed the same amount of K^+^ ions from the corresponding extracellular compartment:

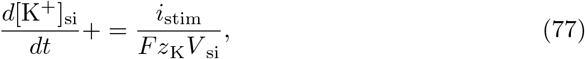

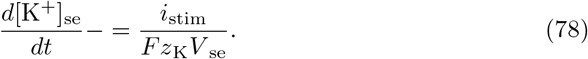

#### Calibration

To let the edPR model calibrate, all simulations were run for 15 s before setting *t* = 0. That is, in all simulations shown in the results sections, the first 15 calibration seconds have been discarded.

#### Analysis

**Fig 9:** To calculate the accumulative transport of ion species k in the intracellular solution (from time zero to *t*) due to diffusion, we integrated *A*_i_*N*_A_*j*_k,diff,i_ from time zero to t, where *N*_A_ is the Avogadro constant. Similarly, we integrated *A*_e_*N*_A_*j*_k,diff,e_ to calculate the accumulative transport of ions in the extracellular solution due to diffusion. We did the same calculations with jk,drift to study the accumulative transport of ions due to drift. When knowing the accumulative transport of each ion species, k_akkum_, we calculated the total transport of e^+^ from their weighted sum:

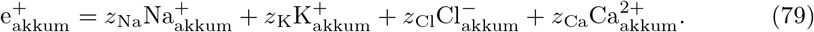

**Fig 10**: To calculate *ϕ*_VC,e_ and *ϕ*_diff,e_, we looked at the extracellular axial current as it is given in the KNP formalism:

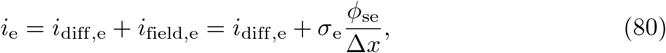

where the last equality follows when we insert Eq 18 for the extracellular field-driven current density *i*_field,e_, and use that *ϕ*_de_ = 0. As in [32], we may split *ϕ*_se_ into two components:

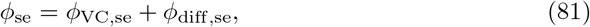

where *ϕ*_vC,se_ is the potential as it would be predicted from standard volume conductor (VC) theory [20, 21], and *ϕ*_diff,se_ is the additional contribution from diffusion [32]. With this, Eq 80 can be written:

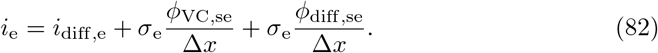

We may split this into two equations if we recognize that

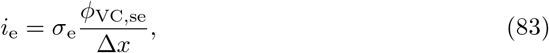

is the standard formula used in VC theory, which is based on the assumption that the extracellular current is exclusively due to a drop in the extracellular VC-potential *ϕ*_VC,se_. The remainder of Eq 82 then leaves us with

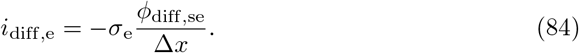

Since we already knew *i*_e_ and *i*_diff,e_ from simulations on the KNP framework, we used Eqs 83 and 84 to calculate *ϕ*_VC,se_ and *ϕ*_diff,se_ separately.

#### Numerical implementation

We implemented the differential equations in Python 3.6 and solved them using the `solve_ivp` function from SciPy. We used its default Runge-Kutta method of order 5(4), and set the maximal allowed step size to 10^-4^. The code is made available at https://github.com/CINPLA/edPRmodel and https://github.com/CINPLA/edPRmodel_analysis.

## References

1. Hodgkin AL, Huxley AF. A quantitative description of membrane current and its application to conduction and excitation in nerve. J Physiol. 1952;117:500–544. doi:10.1007/BF02459568.

2. Traub RD, Wong RK, Miles R, Michelson H. A model of a CA3 hippocampal pyramidal neuron incorporating voltage-clamp data on intrinsic conductances. Journal of Neurophysiology. 1991;66(2):635–650.

3. Pinsky PF, Rinzel J. Intrinsic and network rhythmogenesis in a reduced Traub model for CA3 neurons. Journal of computational neuroscience. 1994;1(1-21):39–60.

4. Mainen ZF, Joerges J, Huguenard JR, Sejnowski TJ. A model of spike initiation in neocortical pyramidal neurons. Neuron. 1995;15(6):1427–1439.

5. Pospischil M, Toledo-Rodriguez M, Monier C, Piwkowska Z, Bal T, Fr?gnac Y, et al. Minimal Hodgkin-Huxley type models for different classes of cortical and thalamic neurons. Biol Cybern. 2008;99(4-5):427–441. doi:10.1007/s00422-008-0263-8.

6. Halnes G, Augustinaite S, Heggelund P, Einevoll GT, Migliore M. A multi-compartment model for interneurons in the dorsal lateral geniculate nucleus. PLoS Comput Biol. 2011;7(9):e1002160. doi:10.1371/journal.pcbi.1002160.

7. Hay E, Hill S, Schürmann F, Markram H, Segev I. Models of neocortical layer 5b pyramidal cells capturing a wide range of dendritic and perisomatic active properties. PLoS computational biology. 2011;7(7):e1002107. doi:10.1371/journal.pcbi.1002107.

8. Sterratt D, Graham B, Gillies A, Willshaw D. Principles of computational modelling in neuroscience. Cambridge University Press; 2011.

9. Offner FF. Ion flow through membranes and the resting potential of cells. The Journal of membrane biology. 1991;123(2):171–182.

10. Koch C. Biophysics of computation: information processing in single neurons. Oxford university press; 2004.

11. Rall W. Core conductor theory and cable properties of neurons. Comprehensive physiology. 2011; p. 39–97.

12. Tveito A, Jæger KH, Lines GT, Paszkowski L, Sundnes J, Edwards AG, et al. An Evaluation of the Accuracy of Classical Models for Computing the Membrane Potential and Extracellular Potential for Neurons. Frontiers in Computational Neuroscience. 2017;11(April):1–18. doi:10.3389/fncom.2017.00027.

13. Traub RD, Contreras D, Cunningham MO, Murray H, LeBeau FE, Roopun A, et al. Single-column thalamocortical network model exhibiting gamma oscillations, sleep spindles, and epileptogenic bursts. Journal of neurophysiology. 2005;93(4):2194–2232.

14. Markram H, Muller E, Ramaswamy S, Reimann MW, Abdellah M, Sanchez CA, et al. Reconstruction and simulation of neocortical microcircuitry. Cell. 2015;163(2):456–492.

15. Arkhipov A, Gouwens NW, Billeh YN, Gratiy S, Iyer R, Wei Z, et al. Visual physiology of the layer 4 cortical circuit in silico. PLoS computational biology. 2018;14(11):e1006535.

16. Buzsáki G, Anastassiou CA, Koch C. The origin of extracellular fields and currents?EEG, ECoG, LFP and spikes. Nature reviews neuroscience. 2012;13(6):407.

17. Einevoll GT, Kayser C, Logothetis NK, Panzeri S. Modelling and analysis of local field potentials for studying the function of cortical circuits. Nature Reviews Neuroscience. 2013;14(11):770–785.

18. Hagen E, Næss S, Ness TV, Einevoll GT. Multimodal modeling of neural network activity: computing LFP, ECoG, EEG and MEG signals with LFPy 2.0. Frontiers in neuroinformatics. 2018;12:92.

19. Anastassiou CA, Koch C. Ephaptic coupling to endogenous electric field activity: why bother? Current opinion in neurobiology. 2015;31:95–103.

20. Holt GR, Koch C. Electrical interactions via the extracellular potential near cell bodies. Journal of computational neuroscience. 1999;6(2):169–184.

21. Pettersen KH, Einevoll GT. Amplitude variability and extracellular low-pass filtering of neuronal spikes. Biophysical journal. 2008;94(3):784–802. doi:10.1529/biophysj.107.111179.

22. Somjen GG. Mechanisms of Spreading Depression and Hypoxic Spreading Depression-Like Depolarization. Physiol Rev. 2001;81(3):1065–1096.

23. Fröhlich F, Bazhenov M. Potassium dynamics in the epileptic cortex: new insights on an old topic. Neuroscientist. 2008;14(5):422–433.

24. Zandt BJ, ten Haken B, van Putten MJ, Dahlem MA. How does spreading depression spread? Physiology and modeling. Reviews in the Neurosciences. 2015;26(2):183–198.

25. Ayata C, Lauritzen M. Spreading depression, spreading depolarizations, and the cerebral vasculature. Physiological Reviews. 2015;95(3):953–993. doi:10.1152/physrev.00027.2014.

26. Herreras O, Somjen G. Analysis of potential shifts associated with recurrent spreading depression and prolonged unstable spreading depression induced by microdialysis of elevated K+ in. Brain research. 1993;610:283–294.

27. Kager H, Wadman WJ, Somjen GG. Simulated seizures and spreading depression in a neuron model incorporating interstitial space and ion concentrations. Journal of neurophysiology. 2000;84(1):495–512.

28. Barreto E, Cressman JR. Ion concentration dynamics as a mechanism for neuronal bursting. Journal of biological physics. 2011;37(3):361–373.

29. Øyehaug L, Østby I, Lloyd CM, Omholt SW, Einevoll GT. Dependence of spontaneous neuronal firing and depolarisation block on astroglial membrane transport mechanisms. Journal of computational neuroscience. 2012;32(1):147–165.

30. Mori Y. A multidomain model for ionic electrodiffusion and osmosis with an application to cortical spreading depression. Physica D: Nonlinear Phenomena. 2015;308:94–108.

31. Halnes G, Mäki-Marttunen T, Keller D, Pettersen KH, Andreassen OA, Einevoll GT. Effect of ionic diffusion on extracellular potentials in neural tissue. PLoS computational biology. 2016;12(11):e1005193.

32. Solbrå A, Bergersen AW, Van Den Brink J, Malthe-Sørenssen A, Einevoll GT, Halnes G. A Kirchhoff-Nernst-Planck framework for modeling large scale extracellular electrodiffusion surrounding morphologically detailed neurons. PLoS computational biology. 2018;14(10):e1006510.

33. Kneller J, Ramirez RJ, Chartier D, Courtemanche M, Nattel S. Time-dependent transients in an ionically based mathematical model of the canine atrial action potential. American Journal of Physiology-Heart and Circulatory Physiology. 2002;282(4):H1437–H1451.

34. Somjen G, Kager H, Wadman W. Computer simulations of neuron-glia interactions mediated by ion flux. Journal of computational neuroscience. 2008;25(2):349–365.

35. Florence G, Dahlem Ma, Almeida ACG, Bassani JWM, Kurths J. The role of extracellular potassium dynamics in the different stages of ictal bursting and spreading depression: a computational study. Journal of theoretical biology. 2009;258(2):219–28. doi:10.1016/j.jtbi.2009.01.032.

36. Cressman JR, Ullah G, Ziburkus J, Schiff SJ, Barreto E. The influence of sodium and potassium dynamics on excitability, seizures, and the stability of persistent states: I. Single neuron dynamics. Journal of computational neuroscience. 2009;26(2):159–170.

37. Ullah G, Cressman Jr JR, Barreto E, Schiff SJ. The influence of sodium and potassium dynamics on excitability, seizures, and the stability of persistent states: II. Network and glial dynamics. Journal of computational neuroscience. 2009;26(2):171–183.

38. Lee J, Kim SJ. Spectrum measurement of fast optical signal of neural activity in brain tissue and its theoretical origin. Neuroimage. 2010;51(2):713–722.

39. Lee J, Boas DA, Kim SJ. Multiphysics neuron model for cellular volume dynamics. IEEE Transactions on Biomedical Engineering. 2011;58(10):3000–3003.

40. Zandt BJ, ten Haken B, van Dijk JG, van Putten MJ. Neural dynamics during anoxia and the “wave of death”. PLoS One. 2011;6(7):e22127.

41. Hübel N, Schöll E, Dahlem MA. Bistable dynamics underlying excitability of ion homeostasis in neuron models. PLoS computational biology. 2014;10(5):e1003551.

42. Dahlem MA, Schumacher J, Hübel N. Linking a genetic defect in migraine to spreading depression in a computational model. PeerJ. 2014;2:e379.

43. Hübel N, Dahlem MA. Dynamics from seconds to hours in Hodgkin-Huxley model with time-dependent ion concentrations and buffer reservoirs. PLoS computational biology. 2014;10(12):e1003941.

44. Wei Y, Ullah G, Ingram J, Schiff SJ. Oxygen and seizure dynamics: II. Computational modeling. Journal of neurophysiology. 2014;112(2):213–223.

45. Wei Y, Ullah G, Schiff SJ. Unification of Neuronal Spikes, Seizures, and Spreading Depression. Journal of Neuroscience. 2014;34(35):11733–11743.

46. Hübel N, Ullah G. Anions govern cell volume: a case study of relative astrocytic and neuronal swelling in spreading depolarization. PloS one. 2016;11(3):e0147060.

47. Hübel N, Hosseini-Zare MS, Žiburkus J, Ullah G. The role of glutamate in neuronal ion homeostasis: A case study of spreading depolarization. PLoS computational biology. 2017;13(10):e1005804.

48. Kager H, Wadman W, Somjen G. Conditions for the triggering of spreading depression studied with computer simulations. Journal of neurophysiology. 2002;88(5):2700–2712.

49. Cataldo E, Brunelli M, Byrne JH, Av-Ron E, Cai Y, Baxter DA. Computational model of touch sensory cells (T Cells) of the leech: role of the afterhyperpolarization (AHP) in activity-dependent conduction failure. Journal of computational neuroscience. 2005;18(1):5–24.

50. Kager H, Wadman W, Somjen G. Seizure-like afterdischarges simulated in a model neuron. Journal of computational neuroscience. 2007;22(2):105–128.

51. Forrest MD, Wall MJ, Press DA, Feng J. The sodium-potassium pump controls the intrinsic firing of the cerebellar Purkinje neuron. PloS one. 2012;7(12):e51169.

52. Chang JC, Brennan KC, He D, Huang H, Miura RM, Wilson PL, et al. A mathematical model of the metabolic and perfusion effects on cortical spreading depression. PLoS One. 2013;8(8):e70469.

53. Le Masson G, Przedborski S, Abbott L. A computational model of motor neuron degeneration. Neuron. 2014;83(4):975–988.

54. Forrest MD. Simulation of alcohol action upon a detailed Purkinje neuron model and a simpler surrogate model that runs¿ 400 times faster. BMC neuroscience. 2015;16(1):27.

55. Krishnan GP, Filatov G, Shilnikov A, Bazhenov M. Electrogenic properties of the Na+/K+ ATPase control transitions between normal and pathological brain states. Journal of neurophysiology. 2015;113(9):3356–3374.

56. Zylbertal A, Kahan A, Ben-Shaul Y, Yarom Y, Wagner S. Prolonged intracellular Na+ dynamics govern electrical activity in accessory olfactory bulb mitral cells. PLoS biology. 2015;13(12):e1002319.

57. Zylbertal A, Yarom Y, Wagner S. The Slow Dynamics of Intracellular Sodium Concentration Increase the Time Window of Neuronal Integration: A Simulation Study. Frontiers in computational neuroscience. 2017;11:85.

58. Qian N, Sejnowski TJ, Diego S. Biological Cybernetics. 1989;15:1–15.

59. Mori Y, Peskin C. A numerical method for cellular electrophysiology based on the electrodiffusion equations with internal boundary conditions at membranes. Communications in Applied Mathematics and Computational Science. 2009;4.1:85–134.

60. Halnes G, Østby I, Pettersen KH, Omholt SW, Einevoll GT. Electrodiffusive model for astrocytic and neuronal ion concentration dynamics. PLoS computational biology. 2013;9(12):e1003386.

61. Niederer S. Regulation of ion gradients across myocardial ischemic border zones: a biophysical modelling analysis. PloS one. 2013;8(4):e60323.

62. Ellingsrud A, Solbrå A, Einevoll G, Halnes G, Rognes M. Finite element simulation of ionicelectrodiffusion in cellular geometries. arXiv preprint arXiv:191103211. 2019;.

63. Attwell D, Laughlin SB. An energy budget for signaling in the grey matter of the brain. Journal of Cerebral Blood Flow & Metabolism. 2001;21(10):1133–1145.

64. Yi G, Fan Y, Wang J. Metabolic cost of dendritic Ca2+ action potentials in layer 5 pyramidal neurons. Frontiers in neuroscience. 2019;13.

65. Tennøe S, Halnes G, Einevoll GT. Uncertainpy: A Python toolbox for uncertainty quantification and sensitivity analysis in computational neuroscience. Frontiers in neuroinformatics. 2018;12.

66. In: Binder MD, Hirokawa N, Windhorst U, editors. Depolarization Block. Berlin, Heidelberg: Springer Berlin Heidelberg; 2009. p. 943–944. Available from: https://doi.org/10.1007/978-3-540-29678-2_1453.

67. Mizuseki K, Royer S, Diba K, Buzsáki G. Activity dynamics and behavioral correlates of CA3 and CA1 hippocampal pyramidal neurons. Hippocampus. 2012;22(8):1659–1680.

68. Gardner-Medwin A. Analysis of potassium dynamics in mammalian brain tissue. The Journal of physiology. 1983;335:393–426.

69. Reiffurth C, Alam M, Zahedi-Khorasani M, Major S, Dreier JP. Na+/K+-ATPase *α* isoform deficiency results in distinct spreading depolarization phenotypes. Journal of Cerebral Blood Flow & Metabolism. 2019; p. 0271678X19833757.

70. Nelson P. Biological physics. WH Freeman New York; 2004.

71. Van Rijn CM, Krijnen H, Menting-Hermeling S, Coenen AM. Decapitation in rats: latency to unconsciousness and the ?wave of death? PloS one. 2011;6(1):e16514.

72. McDougal RA, Hines ML, Lytton WW. Reaction-diffusion in the NEURON simulator. Frontiers in neuroinformatics. 2013;7:28.

73. Hines M, Davison AP, Muller E. NEURON and Python. Frontiers in neuroinformatics. 2009;3:1.

74. Halnes G, Mäki-Marttunen T, Pettersen KH, Andreassen OA, Einevoll GT. Ion diffusion may introduce spurious current sources in current-source density (CSD) analysis. Journal of Neurophysiology. 2017;118(1):114–120. doi:10.1152/jn.00976.2016.

75. Gratiy SL, Halnes G, Denman D, Hawrylycz MJ, Koch C, Einevoll GT, et al. From Maxwell’s equations to the theory of current-source density analysis. European Journal of Neuroscience. 2017;45(8):1013–1023. doi:10.1111/ejn.13534.

76. Rall W. Core conductor theory and cable properties of neurons. In: Kandel ER, Brookhardt JM, Mountcastle V M, editors. Handbook of Physiology. Bethesda: American Physiological Society; 1977. p. 39–97. Available from: http://onlinelibrary.wiley.com/doi/10.1002/cphy.cp010103/full.

77. O’Connell R, Mori Y. Effects of Glia in a Triphasic Continuum Model of Cortical Spreading Depression. Bulletin of Mathematical Biology. 2016;78(10):1943–1967. doi:10.1007/s11538-016-0206-9.

78. Tuttle A, Riera Diaz J, Mori Y. A computational study on the role of glutamate and NMDA receptors on cortical spreading depression using a multidomain electrodiffusion model. PLoS Computational Biology. 2019;15(12).

79. Eisenberg RS, Johnson EA. Three-dimensional electrical field problems in physiology. Progress in biophysics and molecular biology. 1970;20:1–65.

80. Henriquez CS. Simulating the electrical behavior of cardiac tissue using the bidomain model. Critical reviews in biomedical engineering. 1993;21(1):1–77.

81. Sundnes J, Nielsen BF, Mardal KA, Cai X, Lines GT, Tveito A. On the computational complexity of the bidomain and the monodomain models of electrophysiology. Annals of biomedical engineering. 2006;34(7):1088–1097.

82. Lyshevski SE. Nano and molecular electronics handbook. CRC Press; 2016.

83. Chen KC, Nicholson C. Spatial buffering of potassium ions in brain extracellular space. Biophysical journal. 2000;78(6):2776–2797.

84. Errata, Principles of computational modelling in neuroscience;. http://www.compneuroprinciples.org/errata.

85. Wolf JA, Moyer JT, Lazarewicz MT, Contreras D, Benoit-Marand M, O’Donnell P, et al. NMDA/AMPA ratio impacts state transitions and entrainment to oscillations in a computational model of the nucleus accumbens medium spiny projection neuron. Journal of Neuroscience. 2005;25(40):9080–9095.

